# All of Us diversity and scale improve polygenic prediction contextually with greatest improvements for under-represented populations

**DOI:** 10.1101/2024.08.06.606846

**Authors:** Kristin Tsuo, Zhuozheng Shi, Tian Ge, Ravi Mandla, Kangcheng Hou, Yi Ding, Bogdan Pasaniuc, Ying Wang, Alicia R. Martin

## Abstract

Recent studies have demonstrated that polygenic risk scores (PRS) trained on multi-ancestry data can improve prediction accuracy in groups historically underrepresented in genomic studies, but the availability of linked health and genetic data from large-scale diverse cohorts representative of a wide spectrum of human diversity remains limited. To address this need, the All of Us research program (AoU) generated whole-genome sequences of 245,388 individuals (release v7) who collectively reflect the diversity of the USA. Leveraging this resource and another widely-used population-scale biobank, the UK Biobank (UKB) with a half million participants, we developed PRS trained on multi-ancestry and multi-biobank data with up to ∼750,000 participants for 32 common, complex traits and diseases across a range of genetic architectures. We then evaluated effects of ancestry, PRS methodology, and genetic architecture on PRS accuracy across a held out subset of ancestrally diverse AoU participants. Overall, we found that the increased diversity of AoU significantly improved PRS performance in some participants in AoU, especially underrepresented individuals, across multiple phenotypes. Notably, maximizing sample size by combining discovery data across AoU and UKB is not the optimal approach for predicting some phenotypes particularly in African ancestry populations; rather, using data from only AoU for these traits resulted in the greatest accuracy. This was especially true for less polygenic traits with large ancestry-enriched effects, and larger heritability estimates in African ancestry populations, such as neutrophil count (*R*^2^: 0.055 vs. 0.035 using AoU vs. cross-biobank meta-analysis, respectively, because of e.g. *DARC*). Lastly, we calculated individual-level PRS accuracies rather than grouping by continental ancestry, a critical step towards interpretability in precision medicine. Individualized PRS accuracy decays linearly as a function of ancestry divergence, but the slope was smaller using multi-ancestry GWAS compared to using European GWAS. Our results highlight the potential of biobanks with more balanced representations of human diversity to facilitate more accurate PRS for the individuals least represented in genomic studies.

## Introduction

Population-scale biobanks with linked health records and genetic data have enabled an exponential increase in genome-wide association studies (GWAS), significantly expanding our understanding of the genetic basis of diseases^1,2^. Polygenic risk scores (PRS), which aggregate variant-disease associations discovered by GWAS, have been developed for many diseases and traits^3^. For some common, complex diseases, PRS have shown potential in aiding population risk stratification and screening, and their clinical implementation is on the horizon^4–7^. However, the vast majority of data used for GWAS still come from European ancestry (EUR) populations, resulting in the limited transferability of most PRS models to populations of other genetic ancestries^8^. This widely-recognized problem represents one of the most pressing challenges facing the clinical translation of PRS.

Several approaches can help mitigate this critical limitation. Statistical methods that leverage GWAS from multiple populations, including PRS-CSx and others, have been developed^9–12^. Benchmarking studies have evaluated these methods across traits of different genetic architectures using various study designs^13–15^. Complementing these empirical evaluations, theoretical studies have compared observed versus expected accuracies of PRS^13,16–18^. They find that while these methods can improve accuracy in some circumstances, the most direct path to increasing accuracy is through larger and more diverse study populations in GWAS.

Efforts like the Pan-UK Biobank (UKB) Project have maximized usage of current existing data resources by conducting GWAS for thousands of phenotypes using data from multiple ancestry groups, but its ancestral diversity is limited^19^. Other GWAS initiatives like the Global Biobank Meta-analysis Initiative (GBMI)^20^ and disease- and trait-specific consortia, such as the Type 2 Diabetes Global Genomics Initiative^21^ and the Genetic Investigation of ANthropometric Traits (GIANT)^22^, focus on collecting ancestrally diverse data for meta-analysis. The Million Veterans Program is very large and diverse, and has recently conducted pan-trait and -ancestry GWAS, although access to summary statistics is more restricted^23^.

Recent efforts have further expanded the availability of multi-ancestry genomic data. The All of Us Research Program (AoU), launched in 2018 by the National Institutes of Health of the United States, aims to gather health data from at least 1 million participants from diverse backgrounds. As of this study, it has released linked phenotypic and whole-genome sequencing data from 245,388 participants^24^. AoU is one of the largest and most accessible resources of populations traditionally underrepresented in biomedical research, with concerted efforts to capture ancestral diversity^24^. Given the ongoing efforts to increase the diversity of genomic studies, understanding how to best leverage multiple biobank resources to optimally predict complex traits with PRS will be a critical step towards their equitable applications.

Multi-ancestry PRS have been developed for a range of diseases and traits, and these studies have largely concluded that increasing diversity in GWAS substantially improves the transferability of PRS^13,22,25–29^. Recent studies have started utilizing the multi-ancestry data available in AoU^7,30–32^. However, the optimal approach for developing PRS from multi-ancestry studies with large numbers of ancestrally diverse participants across population-scale biobanks remains unclear, especially across traits spanning a range of genetic architectures. Previous studies on optimal strategies for constructing multi-ancestry PRS have mostly used the UKB, which is not fully representative of the broader UK population and has limited ancestral diversity^15,33,34^. Additionally, studies investigating factors contributing to low PRS generalizability have largely focused on phenomena in population genetics, like the outsized impact of differences in allele frequencies and patterns of linkage disequilibrium (LD) on PRS accuracy^16,33^. Yet, there is also clear context-specificity to PRS accuracy that reflects factors like sex-specific heritability differences^35,36^ and biobank-specific characteristics^37,38^. Our understanding of how differences between biobanks – for example, in ascertainment, data collection approaches, and sample recruiting strategy – impact polygenic prediction is still relatively limited. Some work on PRS development using multi-biobank data suggests that increases in sample size from combining heterogeneous biobanks can improve prediction performance for some diseases^25^. Furthermore, recent guidance on individualizing PRS performance evaluations have been based on single ancestry discovery cohorts^39^, and understanding how this applies in multi-ancestry GWAS is an important outstanding question.

In this study, we developed PRS using multi-ancestry and multi-biobank data from AoU and UKB for dozens of commonly-studied diseases and quantitative traits with different genetic architectures. Specifically, we constructed PRS using single-ancestry GWAS from AoU, as well as multi-ancestry meta-analyses within and across AoU and UKB, to investigate the impacts of ancestry composition, sample size, trait genetic architecture, and biobank heterogeneity on PRS accuracy. Given the widespread adoption of UKB data, we also benchmark PRS performance with UKB. We illustrate nuance in optimal PRS strategies across phenotypes, particularly in underrepresented ancestry groups, providing guidelines and reference points for future PRS models developed in diverse genetic studies.

## Results

### Overview of study design

Few frameworks have been developed for analyzing the wealth of phenotypic data available in AoU. We therefore adapted insights from previous UKB analyses. The Pan-UKB Project’s quality control framework, which prioritizes phenotypes based on heritability estimates and other quality metrics, guided our phenotype selection^19^. From these prioritized phenotypes, we selected 14 quantitative and 18 binary phenotypes for our study based on data availability in AoU and other factors (**Methods**, **Supplementary Table 1**).

We assigned participants in AoU to genetically inferred ancestry groups based on principal component analysis (PCA) comparisons with population genetic reference panels (**Methods**). We trained a random forest model using labels from the Human Genome Diversity Panel (HGDP) and 1000 Genomes Project, which we use throughout this study to refer to individuals with genetic ancestry most similar to those in the reference panels: EUR (European), AFR (African), AMR (Admixed American), CSA (Central/South Asian), and EAS (East Asian).

We conducted single-ancestry GWAS in AoU for all phenotypes using data from three groups with the largest sample sizes (N >10,000 including AFR, AMR, and EUR) (**Supplementary Table 1**). We combined GWAS across ancestries through inverse variance-weighted meta-analyses. For comparison, we included discovery GWAS from EUR and AFR populations in the UKB, excluding AMR due small sample size and unreliable genetic association results (**Figure 1**). Finally, we conducted cross-biobank, multi-ancestry meta-analyses.

**Figure 1.**
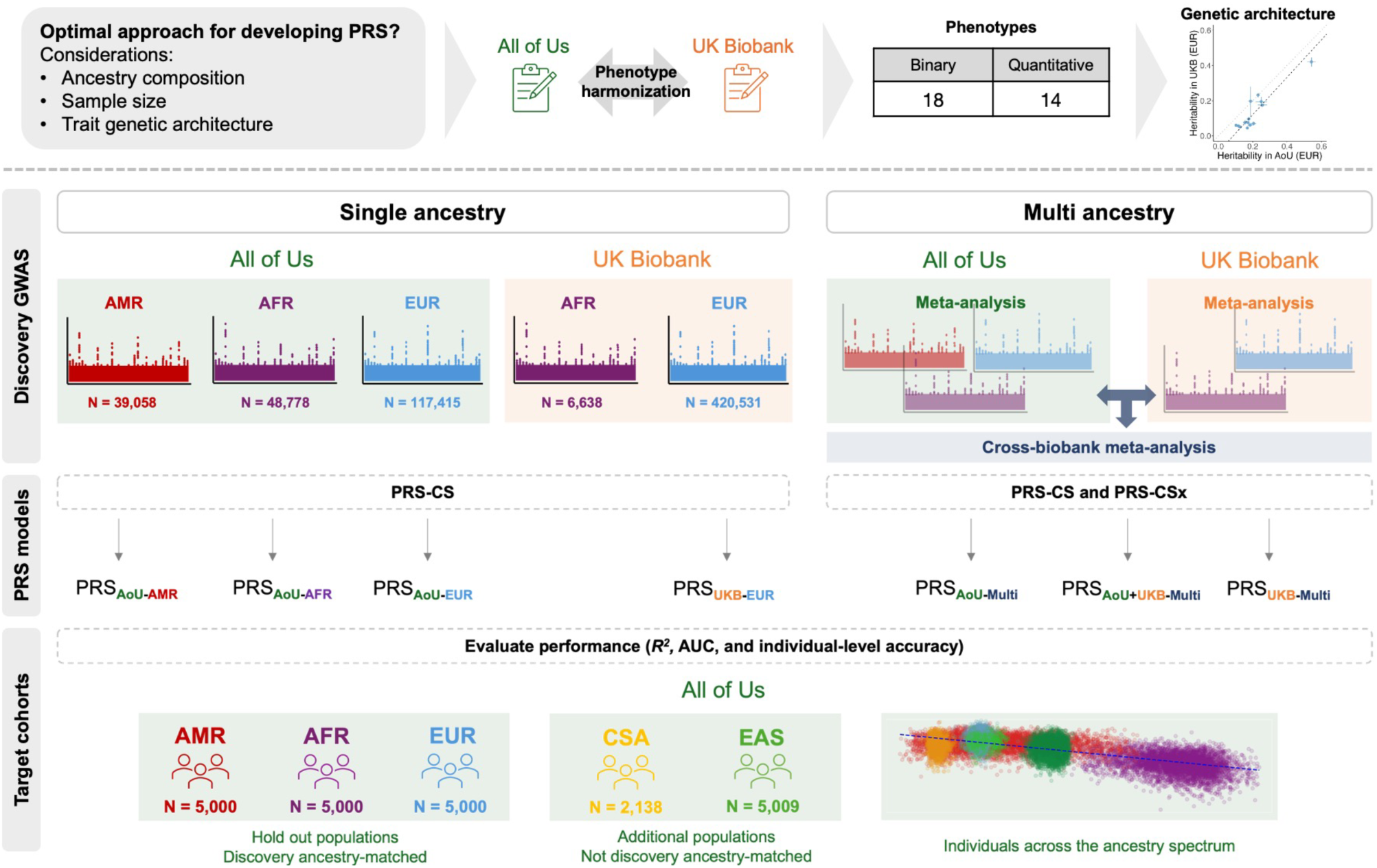
Study design for evaluating optimal PRS strategies that integrate ancestries and biobanks across multiple traits. Overview of workflow showing GWAS used for discovery data, methods for PRS construction, and cohorts used for PRS evaluation. AFR, African; AMR, admixed American; EAS, East Asian; MID, Middle Eastern; EUR, European; CSA, Central and South Asian.

To ensure consistency in phenotype definitions between AoU and UKB, we computed heritability estimates using LD score regression (LDSC)^40^ and genomic relatedness-based restricted maximum likelihood (GREML) as implemented in GCTA^41,42^, as well as genetic correlations across biobanks and population groups using LDSC and Popcorn^43^ (**Methods, Supplementary Tables 2 and 5**). We also compared effect sizes of genome-wide significant associations from biobank-specific GWAS (**Supplementary Fig. 1**) and raw phenotype distributions (**Supplementary Fig. 2**). Overall, our analyses indicated reasonable consistency between AoU and UKB phenotypes^17,44^.

Using these GWAS and meta-analyses as training data, we constructed PRS using two Bayesian, genome-wide methods, PRS-continuous shrinkage (PRS-CS) and its multi-ancestry extension, PRS-CSx, as well as the classic pruning and thresholding method (P + T). We denoted PRS using the following nomenclature: PRS_[biobank]-[ancestry]_, which indicates the GWAS data used to develop the PRS (e.g. PRS_AoU-AFR_ refers to PRS from the GWAS of AFR individuals in AoU); PRS_[biobank]-Multi_ was trained on the multi-ancestry meta-analyses from one or both biobanks (e.g. PRS_AoU+UKB-Multi_ refers to PRS from the meta-analysis of GWAS from multiple ancestries in AoU and UKB). We assessed the performance of each PRS using incremental *R^2^* for quantitative traits and area under the receiver-operating characteristic curve (AUC) for binary phenotypes in five ancestry groups with independent AoU target data (**Methods**). These included unrelated individuals from withheld EUR, AMR, and AFR groups (N=5,000 from each group), as well as CSA (N=2,138) and EAS (N=5,009).

### Target ancestry-matched GWAS improve PRS performance for underrepresented ancestry groups

Although the EUR group is still the largest single ancestry group in AoU, the sample sizes of the AFR and AMR groups in AoU are significantly larger compared to UKB (more than 7 and nearly 40 times larger, respectively). To determine if this increase in sample sizes improves PRS prediction accuracy in underrepresented ancestry groups, we first evaluated PRS constructed from single-ancestry GWAS in AoU. We focused on the results from PRS-CS in the following sections as PRS derived from PRS-CS outperformed or performed comparably to P+T (**Supplementary Tables 9 and 10**), consistent with previous findings^25^. Additionally, a comparison of the relative accuracies of PRS_AoU-EUR_ derived from P+T vs. PRS-CS showed similar transferability of the two methods across the target populations (**Supplementary Fig. 3**). As expected, in the EUR target group, PRS_AoU-EUR_ significantly outperformed PRS_AoU-AFR_ and PRS_AoU-AMR_ across all quantitative traits (median *R^2^*: 0.01 vs. 0.001 and 0.002, Wilcoxon rank sum exact test, p = 6.7e-06 and 5.3e-03, respectively) (**Fig. 2; Supplementary Fig. 4; Supplementary Table 3**). For the AFR and AMR groups, ancestry-matched discovery GWAS often performed best. PRS_AoU-AFR_ achieved the highest median *R^2^ (0.007) i*n the AFR target group across quantitative traits. Similarly, PRS_AoU-AMR_ had highest accuracy in the AMR target group (median R2: 0.01). This indicates that target ancestry-matched discovery GWAS can outperform larger-scale EUR-derived PRS in underrepresented ancestries with the sample sizes currently available in AoU. In the CSA and EAS target groups, PRS_AoU-EUR_ generally performed best, but the median *R^2^* of PRS_AoU-EUR_ in these groups was lower than the the median *R^2^* of the corresponding ancestry-matched PRS in the AFR and AMR target groups, further highlighting the importance of ancestry matching between discovery and target groups. In a sensitivity analysis, we downsampled each of the training datasets in AoU to equal sample sizes to more rigorously evaluate the impact of ancestry matching (**Methods**). The results were consistent: PRS constructed from ancestry-matched, single-ancestry GWAS demonstrated the best overall performance in the EUR, AFR, and AMR target groups (**Supplementary Fig. 5**).

**Figure 2.**
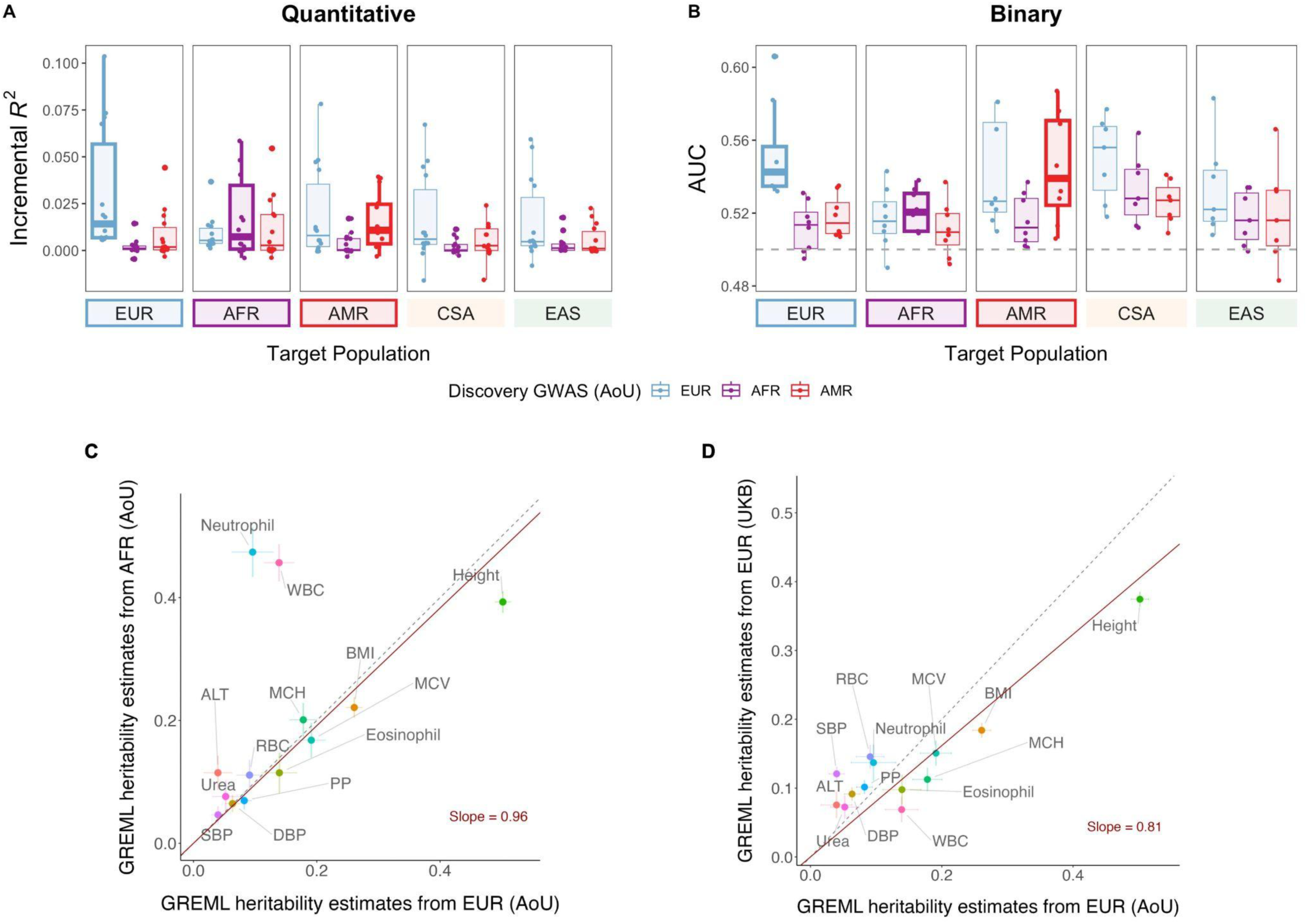
Single-ancestry discovery GWAS from AoU improve PRS performance for ancestry-matched target groups dependent on trait-specific genetic architecture. Each point represents a phenotype, with PRS constructed from PRS-CS reported here, for A) quantitative traits and B) binary traits. Target populations with ancestry-matched PRS are outlined. C) Population groups in AoU and D) EUR groups in AoU and UKB were downsampled to match the smallest sample size of unrelated individuals across training datasets for each phenotype shown. GREML was applied to the individual-level data from these groups to calculate SNP-based heritability. Regression slopes were computed using the Deming regression method^45^. Dashed lines indicate y=x.

Since the UKB has much larger sample sizes of EUR participants compared to AoU, we next investigated whether single-ancestry UKB training data improves prediction for underrepresented ancestry groups in AoU. Specifically, we evaluated PRS_UKB-EUR_ in the AoU target populations. In the AMR target group, PRS_UKB-EUR_ outperformed PRS_AoU-AMR_ for all quantitative traits except neutrophil count, where PRS_AoU-AMR_ showed a 2-fold improvement over PRS_UKB-EUR_ (*R^2^*: 0.02 vs. 0.01) (**Supplementary Fig. 4; Supplementary Table 3**). However, in the AFR target group, PRS_AoU-AFR_ outperformed PRS_UKB-EUR_ for 4 blood panel traits, and achieved comparable accuracy as PRS_UKB-EUR_ for BMI and RBC count (BMI *R^2^*: 0.16 vs. 0.17 and RBC count *R^2^*: 0.11 vs. 0.13) (**Supplementary Fig. 4**). The >20-fold greater sample size of the EUR UKB vs. AFR AoU discovery groups (N=407,810 vs. N=18,044) did not result in significant PRS performance improvement for these traits. These results highlight the importance of training PRS on discovery cohorts that match the ancestry of target populations, particularly those with significant genetic differentiation from majority populations. Vast increases in EUR discovery sample sizes cannot compensate for the lack of training data from underrepresented groups.

To explore how various factors beyond the well-known differences in LD and minor allele frequency contribute to our findings that target-ancestry matched GWAS can yield better PRS performance even with smaller sample sizes, we further estimated SNP-based heritability, which theoretically bounds PRS accuracy, using LDSC for EUR groups in both UKB and AoU (**Methods, Supplementary Table 5**). We found that the heritability in the AoU was overall lower than that in the UKB, likely reflecting the heterogeneity between biobanks. To account for the impact of sample size, we further estimated heritability using GREML^41,42^ with matched sample sizes of the different discovery ancestry groups within AoU and also the EUR groups in AoU and UKB (Methods). We found that the estimates were highly consistent across the AoU groups for quantitative traits, except for a few blood panel traits with ancestry-enriched variants (**Fig. 2C, Supplementary Fig. 16, Supplementary Table 5**). The estimates were less aligned but largely consistent as well for EUR groups between AoU and UKB (Deming regression slope = 0.81) (**Fig. 2D**), reflecting biobank-specific characteristics affecting heritability estimates.

In the context of cross-ancestry prediction, PRS accuracy is also affected by genetic correlation. Specifically, expected prediction accuracy can be approximated as: 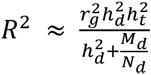, where 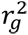 denotes cross-ancestry genetic correlation, 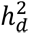 denotes SNP-based heritability in the discovery population, 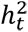 denotes SNP-based heritability in the target population, *M_d_* is the number of independent chromosome segments in the discovery population and *N_d_* is the discovery GWAS sample size^13^. The estimated cross-population genetic correlations using Popcorn (**Methods**) between EUR, AFR, and AMR groups in AoU, were less than 1 for most quantitative traits (**Supplementary Table 2**). Imperfect and smaller genetic correlations likely contributed to the superior performance of target ancestry-matched PRS_AoU_ compared to other single-ancestry PRS_AoU_. Genetic correlation estimates between the EUR groups in AoU and UKB were closer to 1 compared to between populations in AoU for most quantitative traits (**Supplementary Table 2**). Given the relative consistencies between the biobanks, as well as low F_ST_ (0.02) between the AMR group in AoU and the EUR group in UKB indicating relatively low genetic differentiation (**Supplementary Fig. 6**), the much larger sample sizes of PRS_UKB-EUR_ likely contributed to the better performance of PRS_UKB-EUR_ compared PRS_AoU-AMR_ in the AMR target group for all quantitative traits except neutrophil count. Neutrophil count had a substantially higher heritability estimate in matched sample sizes of AoU AMR (0.30, SE = 0.03) vs. UKB EUR (0.14, SE = 0.03), especially compared to the other quantitative traits; additionally, it had the lowest genetic correlation estimate between AoU AMR and AoU EUR (rg = 0.40), which was significantly different from 1. Similarly, the four traits for which PRS_AoU-AFR_ outperformed PRS_UKB-EUR_ (MCH, MCV, WBC, and neutrophil count) had higher heritability estimates in matched sample sizes of AoU AFR compared to UKB EUR. Notably, the genetic distance between AoU AFR and UKB EUR is also higher (F_ST_ = 0.08). Phenotype distributions were mostly similar across the biobanks and populations (**Supplementary Fig. 2**), indicating that sample size and ancestry matching likely had the greatest impacts on PRS performance in this study.

We next investigated PRS performance for the binary phenotypes to compare with the well-powered quantitative traits. Due to overall smaller sample sizes (**Supplementary Table 1**), we limited evaluation of their PRS to diseases with at least 10,000 cases and larger heritability estimates (>0.03 in EUR using LDSC), which included chronic ischaemic heart disease, chronic obstructive pulmonary disease (COPD), asthma, type 2 diabetes (T2D), lipid metabolism disorders, coronary atherosclerosis, esophagitis, and kidney stones. We observed similar patterns in PRS performance across these 8 disorders as we observed for the quantitative traits: the ancestry-matched PRS achieved the highest median AUC in each of the EUR, AFR, and AMR target groups (**Fig. 2**). Of note, in the AFR target group, PRS_AoU-AFR_ achieved the highest AUC for asthma (0.54), comparable to the AUC of PRS_UKB-EUR_ (0.54), as well as COPD (0.53) (**Supplementary Table 4; Supplementary Fig. 7)**. PRS_AoU-AFR_ also had the highest prediction accuracy as measured by variance explained on the liability scale 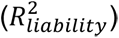 for asthma (0.007), COPD (0.004), disorders of lipid metabolism (0.003), and esophagitis (0.0005) in the AFR group. The prevalences of asthma and COPD in the AoU AFR group (18% and 9%, respectively) are significantly higher than the prevalences in AoU EUR (16% and 8%), potentially contributing to the higher prediction accuracies of PRSAoU-AFR (**Supplementary Table 1**). However, disorders of lipid metabolism and esophagitis had significantly higher prevalences in AoU EUR (50% and 37%) compared to in AoU AFR (35% and 30%). Therefore, disease prevalence may contribute to differences in PRS performance but ancestry matching between discovery and target cohorts may play a greater role.

### Integrating multiple ancestries for discovery GWAS can improve PRS performance compared to single-ancestry GWAS

Building on recommendations from previous studies^13,25,28^, we next investigated how multi-ancestry meta-analyses affect PRS accuracy across quantitative and binary phenotypes. We first evaluated multi-ancestry meta-analyses from the UKB, and found that PRS_UKB-Multi_ showed little to no improvement in PRS performance compared to PRS_UKB-EUR_ across the target groups in AoU due to the vastly different sample sizes between EUR and AFR groups in the UKB (**Supplementary Table 3**).

We then evaluated the performance of PRS derived from the multi-ancestry AoU meta-analyses. Across the quantitative traits, PRS_AoU-Multi_ had comparable accuracy to PRS_AoU-EUR_ and PRS_AoU-AMR_ in the EUR and AMR target groups, respectively (**Supplementary Table 3**). In the AFR target group, we observed an improvement of 0.6% in median *R^2^* compared to PRS_AoU-AFR_, and PRS_AoU-Multi_ outperformed PRS_AoU-AFR_ for all quantitative traits. Accuracy gains from PRS_AoU-Multi_ were especially large for some traits, including body mass index (BMI), mean corpuscular hemoglobin (MCH), mean corpuscular volume (MCV), neutrophil count, and white blood cell (WBC) count. Comparing the AoU and UKB meta-analyses, we found that in the EUR and AMR target groups, PRS_AoU-Multi_ had lower performance across the traits compared to PRS_UKB-Multi_ (**Fig. 3A, Supplementary Fig. 8A**). The EUR group dominates the multi-ancestry UKB meta-analyses, and given that PRS_UKB-EUR_ outperformed the target-ancestry matched PRS in these groups while PRS_AoU-Multi_ did not, the difference in performance between PRS_AoU-Multi_ and PRS_UKB-Multi_ was expected. As noted previously, the low genetic differentiation, measured by *F_ST_*, between the AMR in AoU and EUR in UKB, as well as between the EUR groups in both biobanks, further supports these results (**Supplementary Fig. 6**): not only is the EUR group in the UKB meta-analyses much larger than in the AoU meta-analyses, it is also genetically proximal to the AMR and EUR groups in AoU, thus contributing to the superior performance of PRS_UKB-Multi_.

**Figure 3.**
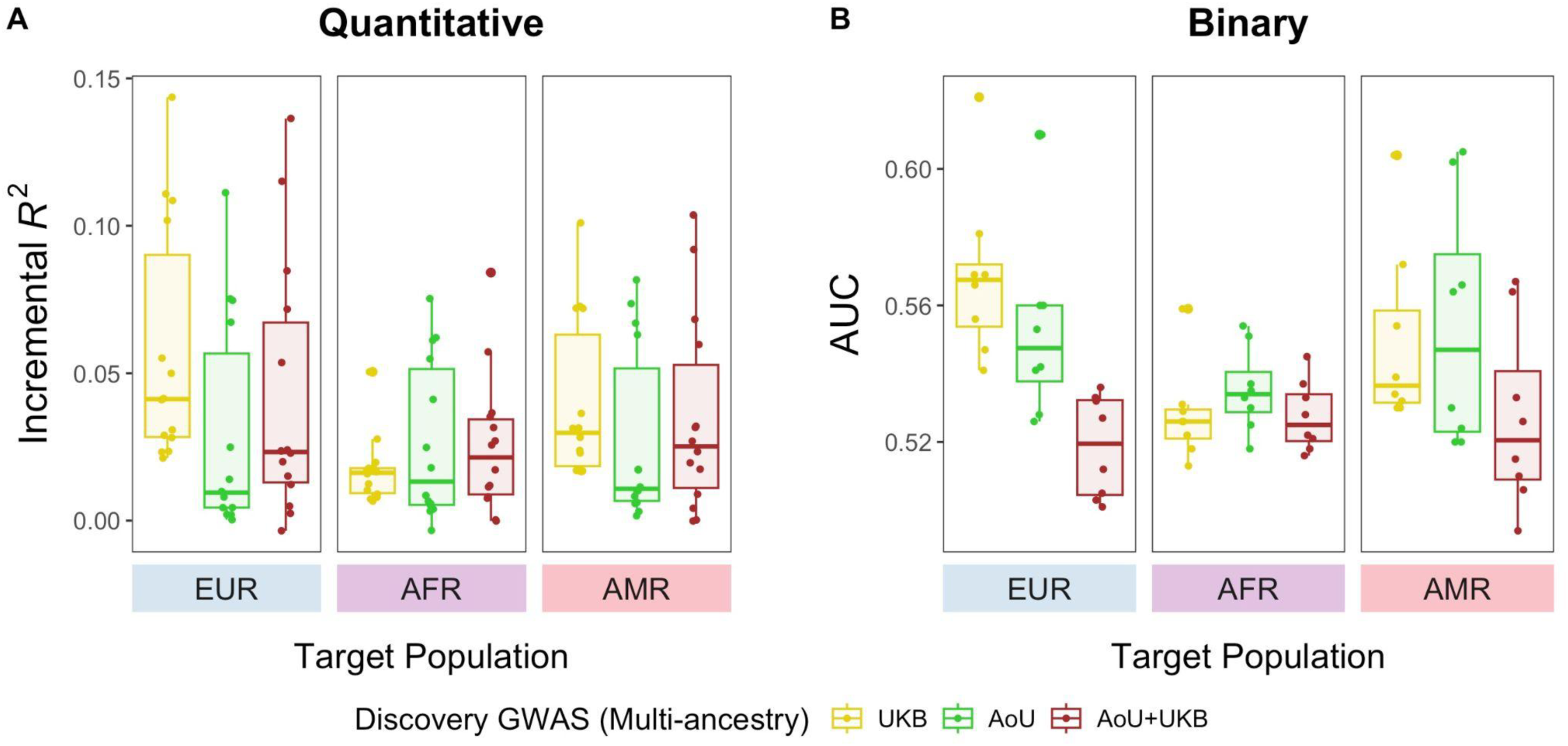
PRS derived from multi-ancestry meta-analyses show variable performance across target groups. Performance of PRS constructed from PRS-CS applied to UKB, AoU, and cross-biobank (AoU and UKB) multi-ancestry meta-analyses are reported here. Each point represents a phenotype.

To gauge the value of combining AoU and UKB for discovery, we next evaluated PRS derived from the cross-biobank multi-ancestry meta-analyses. In the AFR target group, PRS_AoU+UKB-Multi_ offered some improvement in median *R^2^* compared to PRS_AoU-Multi_ and PRS_UKB-Multi_ (0.021 vs. 0.013 and 0.016, respectively) (**Fig. 3A, Supplementary Fig. 8A**). However, that improvement depended on genetic architecture: prediction in more polygenic traits (**Supplementary Table 6**) such as BMI and DBP benefited from the increase in sample size in the cross-biobank meta-analyses; conversely, PRS_AoU-Multi_ outperformed PRS_AoU+UKB-Multi_ for less polygenic traits or those with large-effect ancestry-enriched variants, such as MCH and MCV. Additionally, as mentioned above, both genetic correlation and differences in SNP-based heritability between target and discovery datasets influence cross-ancestry prediction performance. While methods for accurately estimating these values in multi-ancestry discovery data remain limited, our results based on single-ancestry discovery data suggest that lower genetic correlation across biobanks and populations, along with divergent SNP-based heritability, likely contribute to the observation that integrating multiple discovery populations does not always improve prediction performance.

PRS_AoU-Multi_ had varying performance in the disease phenotypes as well (**Supplementary Table 4**). In the AFR target group, PRS_AoU-Multi_ showed increased performance compared to PRS_AoU-AFR_ for a couple diseases (ischaemic heart disease and coronary atherosclerosis) but did not improve prediction performance in diseases where PRS_AoU-AFR_ outperformed PRS_AoU-EUR_ (COPD, asthma, and lipid metabolism disorders). In the AMR target group, PRS_AoU-Multi_ marginally improved AUC compared to PRS_AoU-AMR_ for only a couple diseases as well (T2D and COPD). In both the AFR and AMR target groups, PRS_AoU+UKB-Multi_ did not offer improved prediction compared to PRS_AoU-Multi_ or any single-ancestry PRS_AoU_ across the diseases (**Fig. 3B, Supplementary Fig. 8B**).

Finally, we compared the performances of multi-ancestry PRS developed using PRS-CS vs. PRS-CSx (**Supplementary Table 7; Supplementary Table 8**). In the AFR target group across the quantitative traits, PRS-CSx improved median *R^2^* by 0.008 over PRS-CS for PRS_AoU-Multi_, with substantial improvements in alanine aminotransferase, BMI, MCH, MCV, and red blood cell (RBC) count. PRS-CSx did not significantly improve performance of PRS_AoU-Multi_ in the EUR or AMR target groups. Across the binary phenotypes, applying PRS-CSx did not improve performance of PRS_AoU-Multi_ in the EUR, AFR, and AMR target groups.

### Optimal PRS differs across phenotypes and target ancestries

To identify the best-performing PRS, we compared all PRS models constructed from PRS-CS for each phenotype, focusing on the target groups with ancestry-matched PRS (**Fig. 4; Supplementary Fig. 9**). We tested for significant differences of prediction accuracy between each PRS and PRS_UKB-EUR_, the best-powered single-ancestry PRS in this study (Wald test, p-value < 0.05 indicates significance). . Improvements in PRS accuracy using data from AoU were greatest for the AFR target group: for many of the traits, PRS derived from multi-ancestry AoU data had significantly higher accuracy than PRSUKB-EUR. For four quantitative traits (MCH, MCV, WBC count, and neutrophil count), PRS_AoU-Multi_ offered more than 4-fold increase in accuracy over PRS_UKB-EUR_ in the AFR group. Altogether, this underlines the importance of using target ancestry-matched discovery data especially for populations with large genetic distances from the predominant EUR populations.

**Figure 4.**
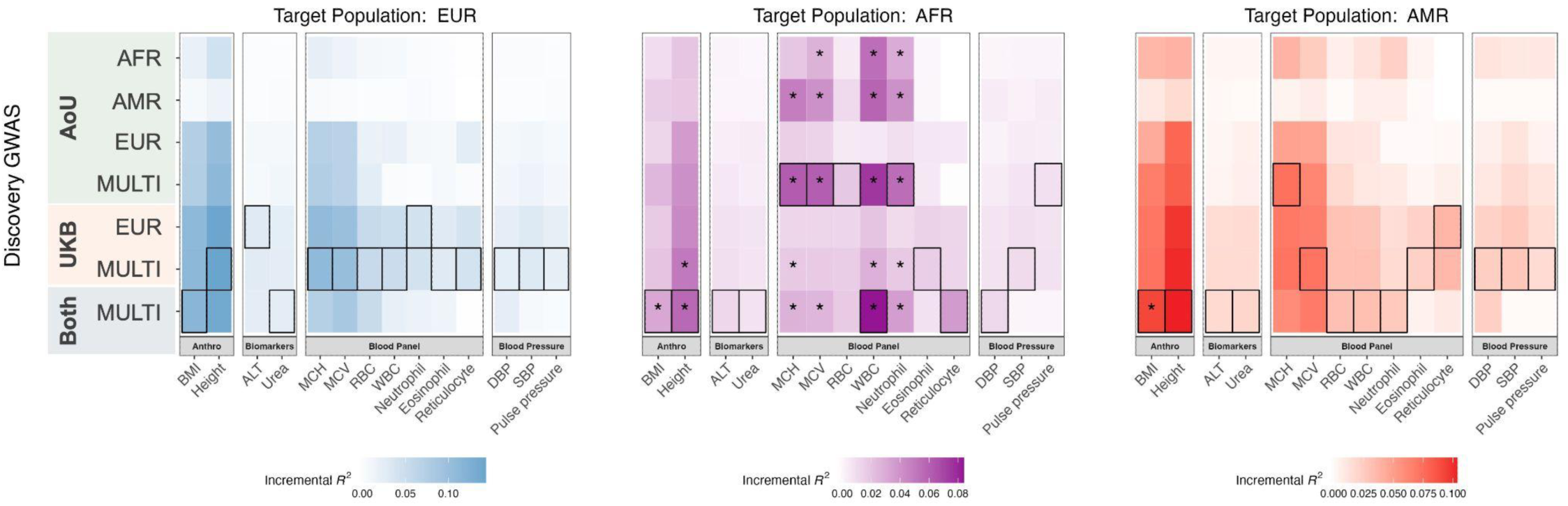
AoU discovery data improve PRS performance in AFR target group. Performance of all PRS models, denoted on y-axis, across quantitative traits, denoted on x-axis. PRS model with greatest *R^2^* per trait is outlined. Asterisk indicates significantly greater prediction accuracy than that of the PRS derived from the EUR UKB discovery group (Wald test, p < 0.05).

Not only does PRS accuracy tend to decay as target populations become more genetically divergent from the training population, but even within populations PRS accuracy can vary because human diversity exists along a continuous spectrum of genetic ancestry rather than in discrete, homogeneous groups. For example, to illustrate this point, we grouped the target populations by proportion of European ancestry rather than genetic ancestry group and found that PRS accuracy increases as the proportion of European ancestry increases when using AoU EUR as the training population (**Supplementary Fig. 10**). Based on recent work proposing a shift from population- to individual-level metrics of PRS accuracy^39,46^, we next explored the impact of individual variation across the genetic ancestry continuum on PRS performance and examined individual-level PRS accuracy as a function of genetic distance (GD) using single- and multi-ancestry AoU discovery data (**Methods**). We focused on the four blood panel traits for which PRS_AoU-Multi_ performed best in the AoU AFR group, as well as two additional well-powered quantitative traits for which PRS_AoU-Multi_ did not outperform PRS_UKB-EUR_ (height and BMI). For baseline comparison, we computed individual PRS accuracy using the EUR GWAS from AoU. Across the blood panel traits, height, and BMI, PRS accuracy decreased with increasing GD from both the EUR and multi-ancestry discovery groups, consistent with previous findings^39^ (**Supplementary Fig. 11**, **Fig. 5A, Supplementary Fig. 13**). Among the blood panel traits, we observed the largest decay in individual-level PRS accuracy in neutrophil count, WBC count, and MCV using the AoU EUR discovery GWAS, described by more negative slopes and lower intercepts (slopes = −2.63, −2.06, and −0.88; intercepts = 0.71, 0.78, and 0.89) (**Fig. 5A, Supplementary Fig. 14**). In contrast, individual-level accuracy computed from the multi-ancestry AoU meta-analyses showed nearly no decay across the genetic ancestry spectrum for neutrophil and WBC count, and slight decay for MCV (slopes = −0.02, −0.01, and −0.42; intercepts = 1.00, 1.00, and 0.94). However, for BMI, individual-level accuracy from the multi-ancestry meta-analysis showed greater decay than the EUR GWAS (slopes = −0.72 vs. −0.63). For MCH and height, the linear decay in individual-level accuracy was still present using the multi-ancestry meta-analyses as discovery, but that decay was attenuated(**Supplementary Fig. 12, Supplementary Fig. 13**).

**Figure 5.**
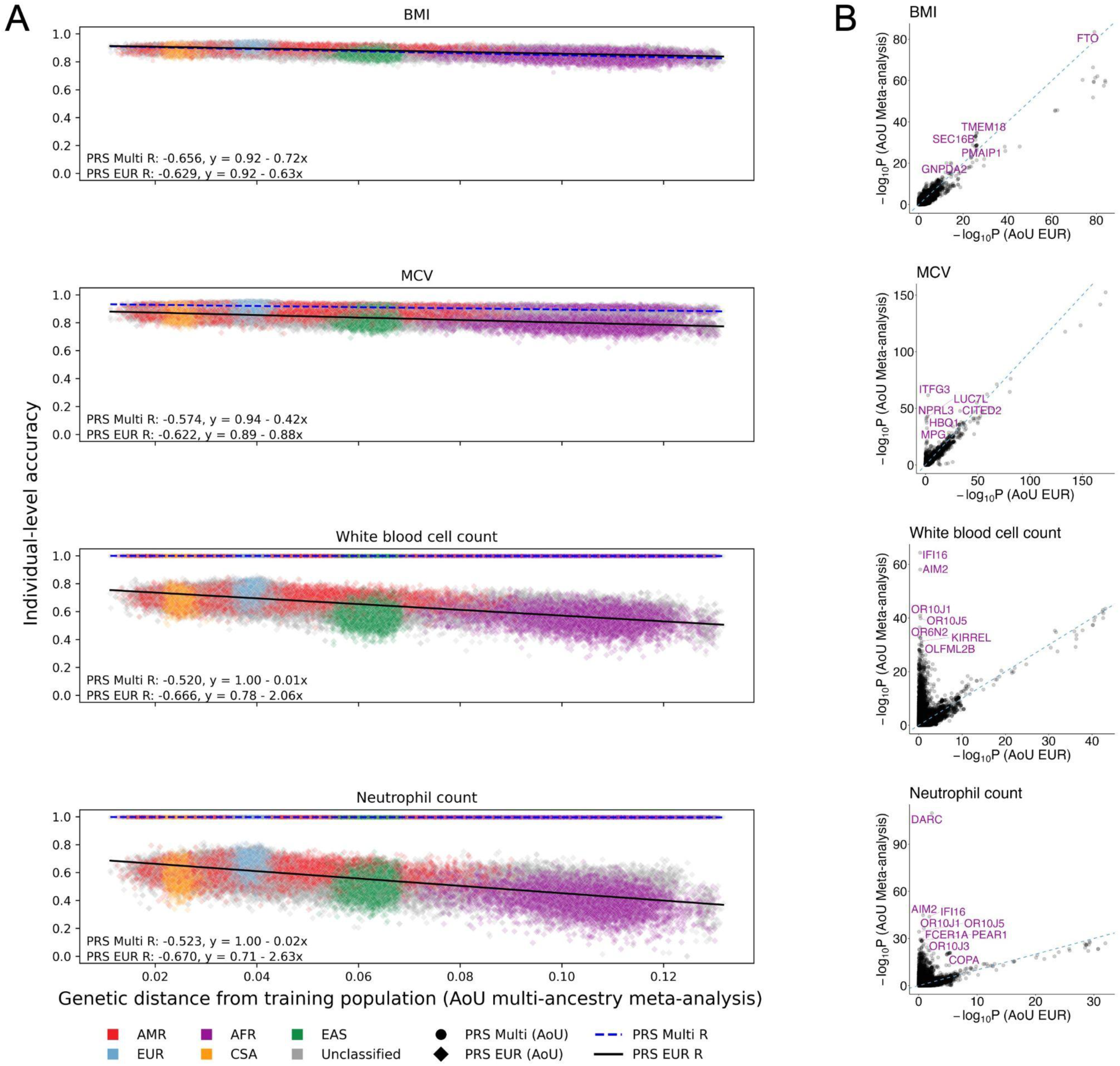
PRS derived from multi-ancestry meta-analyses for blood panel traits show improved accuracy on individual-level, driven by ancestry-enriched variants. A) Individual-level accuracy of PRS derived from AoU multi-ancestry meta-analyses and EUR GWAS across target individuals in AoU, represented by each point. The x-axis represents the genetic distance (GD) of each target individual from the combined discovery populations included in the AoU multi-ancestry meta-analyses. The y-axis shows the PRS accuracy, which was scaled to enable cross-trait comparisons of decay in accuracy as a function of GD; as a result, proportions of genetic liability explained by PRS for each individual are not represented here. *R* was calculated as the correlation between GD and PRS accuracy from a two-sided Pearson correlation test. The colors represent genetic ancestry groups as inferred by PCA. B) Comparison of GWAS significance in AoU multi-ancestry meta-analyses and AoU EUR GWAS across blood panel traits. SNPs tested in both the AoU multi-ancestry meta-analyses and EUR GWAS are represented by each point. SNPs reaching genome-wide significance (p < 5e-8) in the AoU meta-analysis and AoU AFR GWAS for each phenotype are annotated. Dashed lines indicate y=x; x- and y-axis scales are specific to each phenotype and differ according to scale of significance in meta-analyses vs. EUR GWAS.

Studies^33,47^ have previously highlighted that greater diversity in the discovery data showed outsized improvements in PRS accuracy for certain blood panel traits, including MCV and WBC count, like y due to specific genetic loci that disproportionately explain population-specific risk and are more common in underrepresented ancestry groups. Indeed, we found that a few genome-wide significant loci from the AFR GWAS in AoU were highly significant in the AoU meta-analyses but not the EUR GWAS, including those closest to *DARC* associated with neutrophil count and *ITFG3* associated with MCH and MCV (**Fig. 5B**), likely driving the increased accuracy of PRS_AoU-Multi_ in the AFR group and for individuals furthest in GD from the discovery data. While phenotype distributions of these blood panels traits were largely consistent across AoU populations (**Supplementary Fig. 2**), neutrophil count and WBC in particular had much higher heritability estimates in matched sample sizes of AoU AFR compared to AoU EUR (h^2^ = 0.47 vs. 0.10 for neutrophil count; 0.46 vs. 0.14 for WBC) (**Fig. 2C, Supplementary Table 5**). These traits also had relatively lower polygenicity estimates, ranging from 0.011-0.014, compared to the other quantitative traits (**Supplementary Table 6**). Thus, these findings suggest that differences in genetic architecture, as well as population-specific genetic factors, contribute to improved accuracy from AoU multi-ancestry training data on both the population- and individual-level.

## Discussion

PRS are already being tested in clinical settings for a variety of diseases^48^. For example, the eMERGE Network identified, validated, deployed, and returned PRS to patients for 10 clinical conditions, including heart disease, asthma, and type 1 and 2 diabetes^7^. This study ultimately spanned four years, highlighting the challenge of translating rapidly evolving GWAS findings into clinical practice. The extensive polygenicity of common complex diseases and the accelerating pace of GWAS discoveries for most diseases complicates timely clinical translation^1^. Nimbleness is needed for PRS to be maximally effective in the clinic. However, studies have shown poor agreement between individuals at the extremes of the PRS distribution when using different GWAS with a best-case overlap of 60% of individuals above the 80th percentile^49^. Additionally, while it is widely recognized that PRS have different accuracies across ancestry groups mostly due to LD and allele frequency differences^8^, PRS generalizability remains a critical challenge; large-scale datasets most commonly used for PRS development and evaluation are often skewed in representation.

The AoU offers a substantially more diverse resource of phenotypic and genomic data compared to other large-scale contemporary biobanks. This important step towards diversifying human genetic datasets raises new questions for PRS development, particularly for historically underrepresented groups. Our study investigated whether the sample sizes of diverse ancestry groups currently available in AoU are sufficient to increase PRS performance. We found that individuals in the AFR target group benefited most from AoU data, particularly from multi-ancestry meta-analyses. However, AoU discovery data did not significantly improve PRS accuracy in other ancestry groups compared to the largest EUR GWAS from UKB. Encouragingly, for some traits with ancestry-enriched variants, AoU multi-ancestry meta-analyses substantially improved PRS accuracy for individuals with genetic distances furthest from the training data for quantitative traits. Sample sizes of disease outcomes are currently somewhat limited across ancestry groups in AoU, making it difficult to discern clear patterns in PRS development for binary traits. For example, although the target ancestry-matched PRS from AoU outperformed non-ancestry-matched PRS across several diseases, stratifying the AMR and AFR target populations based on the target ancestry-matched PRS slightly improved risk stratification for only a few diseases (Supplementary Fig. 15).

Combining AoU and UKB GWAS in cross-biobank meta-analyses did not uniformly improve accuracy across the phenotypes and target groups, despite the increase in sample size. Across ancestries and traits, the optimal PRS in AoU target populations tended to be the largest cross-biobank multi-ancestry meta-analysis for EUR, the largest meta-analysis or the UKB multi-ancestry meta-analysis for AMR (excluding AoU data), and the largest meta-analysis or the AoU multi-ancestry meta-analysis for AFR (excluding UKB data). These observations highlight the complexity of fine-tuning and improving PRS accuracy, which is affected by factors beyond solely sample size. Additional factors include genetic divergence, heritability, and the degree of phenotype consistency. Heritability estimates in our study using matched sample sizes and individual-level data were relatively consistent between the EUR groups across biobanks for quantitative traits. When using the full sample sizes of imbalanced data and a summary-statistics based method, heritability estimates were higher in UKB, likely due to more precise effect size estimates in the larger study. Sample size-matched heritability estimates were largely consistent between populations within AoU, with the exception of some traits with large ancestry-enriched variants that are not identified in EUR GWAS alone; phenotype distributions across ancestry groups and biobanks were also mostly consistent. On the other hand, genetic correlation estimates were on average lower between the EUR and the AMR and AFR groups in AoU compared to between the EUR groups across AoU and UKB. Together, these observations may help explain the relative importance of ancestry-matching over solely increasing sample size for some traits. Nevertheless, no single strategy is appropriate for all scenarios. Compounding these complexities, while differences in disease prevalence across populations and biobanks have typically identified consistent genetic effects, subsequent context-specific calibration is critical to maximize translational utility^20,48^. Understanding the impacts of inter- and intra-biobank heterogeneity on PRS accuracy will be important as AoU and other biobanks, like the Million Veteran Program^23^, continue to grow in scale and diversity.

As the trajectory of PRS development advances towards clinical implementation, understanding the absolute risk conferred by PRS is crucial for translation. Although individualizing PRS metrics of accuracy is an important step towards translation, additional investigations into the calibration and interpretation of PRS will be needed. For example, integrating PRS into clinical models with other known risk factors that vary in frequency across healthcare systems is an important area for future investigation. Future work should also assess the effects of non-genetic risk factors, which differ across individuals and populations, on PRS accuracy as more clinical and environmental data becomes available in AoU and other diverse biobanks.

## Methods

### Datasets and quality control

#### Pan-UK Biobank (Pan-UKB)

The UK Biobank (UKB) is an extensively utilized cohort comprising approximately 500,000 participants from the United Kingdom, ranging in age from 40 to 69 years. Detailed documentation concerning this cohort has been previously reported^50^. In pursuit of harnessing the rich diversity present within the UKB beyond the customary European ancestry individuals, the Pan-UKB project (https://pan.ukbb.broadinstitute.org/) has undertaken a comprehensive multi-ancestry investigation. This project encompasses 7,228 distinct phenotypes across 6 continental ancestry groups, with a cumulative total of 16,131 GWAS. Rigorous quality control procedures were applied to scrutinize the phenotypic-level, individual-level, and variant-level data, with comprehensive details available in Karczewski et al.^19^

#### The All of Us Research Program (AoU)

The All of Us Research Program, launched by National Institute Health in May 2018, represents a longitudinal cohort study with the goal of engaging at least 1 million participants encompassing diverse ancestral backgrounds. By leveraging comprehensive data collection including biospecimens, health questionnaires, electronic health records and physical measurements, AoU aims to advance precision medicine and enhance overall human health^51^. Participants, aged 18 years and older, are recruited from over 340 centers with informed consent. As of April 2023, a subset of around 250,000 participants has undergone whole genome sequencing (WGS). We assigned those individuals with WGS data into the nearest genetic ancestry based on principal components (PCs), resulting in 49,778 of African descent (AFR), 39,058 of American descent (AMR), 2,138 of Central and South Asian descent (CSA), 5,183 of East-Asian descent (EAS), 117,415 of European descent (EUR) and 432 of Middle Eastern descent (MID). The strategy was the same as described in the pan-UKB project^19^. Briefly, we projected all AoU individuals into the PC space using pre-estimated weights of 168,899 variants^20^ from the Human Genome Diversity Panel (HGDP)^52^ and 1000 Genomes Project^53^. For individuals with a probability > 50% from the random forest, we further refined initial ancestry assignments by pruning outliers within each continental assignment. We reran PCA within each assigned continental ancestry group and calculated total distances from population centroids across 10 PCs. Using these PC scores, we computed centroid distances across 3-5 centroids based on the heterogeneity within each group. We identified and removed ancestry outliers by plotting histograms of centroid distances and excluding individuals at the extreme high end.

Given the limited sample size within CSA, EAS and MID ancestral populations, we exclusively used them as independent test cohorts. For EUR, AMR and AFR populations, we split the data into separate training and test sets. Specifically, in each population, we randomly selected 5,000 individuals from unrelated samples as the withheld test dataset. We used the remaining individuals as the training dataset, which included related individuals to improve statistical power. To avoid relatedness between test and training dataset, we subsequently removed individuals in the training dataset that showed a kinship coefficient larger than 0.1 with any individual in the test dataset. The estimates of kinship coefficient were provided by AoU. We removed those individuals who did not pass AoU quality controls. Consequently, we used 43,926, 33,330 and 111,850 individuals as the training dataset for AFR, AMR and EUR, respectively. For the variant-level quality controls, we focused on only HapMap 3 variants and further removed those with minor allele frequency (MAF) lower than 0.01, genotype missing rates larger than 0.05 and hardy-weinberg equilibrium (HWE) *p*-value smaller than 1e-6.

### Phenotypes

#### UKB

For those 492 high quality phenotypes that passed different filters as described in Karczewski et al.^19^, we calculated the variance explained by the top genome-wide significant loci as ∑ 2*p*(1 − *p*)*β*^2^ where *p* is the MAF and *β* denotes the estimated per-allele effect sizes on the standardized phenotype. The top loci were defined using clumping in PLINK^54^ based on ancestry-matched reference panels from UKB; more details can be found in Karczewski et al.^19^. We identified a subset of 129 phenotypes, characterized by a greater variance explained in the multi-ancestry meta-analyzed GWAS in comparison to EUR-based GWAS. We focused on this subset of phenotypes, considering the potential to improve predictive accuracy in underrepresented populations by leveraging multi-ancestry discovery GWAS. Subsequently, those selected phenotypes were subject to further in-depth investigation in the AoU.

#### AoU

To enhance the quality and reliability of the phenotypic data available within the AoU, we curated and processed the phenotypes through a few steps. First, we checked whether there are matched phenotype descriptions in AoU based on data-field notes in the UKB showcase (https://biobank.ndph.ox.ac.uk/showcase/). Phenotypes derived from survey data were subsequently excluded from consideration. Following this filtering process, phenotypes with either matched or closely related descriptions in AoU were selected for further evaluation. We also added a few commonly studied quantitative traits (BMI, height, and eosinophil count), as well as three additional common diseases with high impact on public health (COPD, asthma, and coronary atherosclerosis). This resulted in 14 quantitative phenotypes, 7 ICD-10 codes and 11 PheCodes for all downstream analyses (**Supplementary Table 1**). The curation of raw phenotypic data encompassed a comprehensive analysis based on concept IDs, and the most recent measurements were sourced from diverse domains, such as conditions, lab and physical measurements, and surveys. For the PheCode curation, we employed the PheCode map v1.2 (https://phewascatalog.org/phecodes) to map ICD codes into corresponding phecodes. Notably, lab and physical measurements often exhibited variations in measurement units across individuals. To address this issue, the most frequent unit of measurement was adopted as a reference, and appropriate conversions were applied to standardize other units accordingly. In order to optimize the sample size available for analysis, individuals for whom the unit concept name was indicated as “empty”, "no matching concept," or "no value" were retained in the dataset. For quantitative phenotypes, individuals with values exceeding 5 standard deviations from the mean were systematically excluded from the dataset to ensure the robustness of subsequent analyses.

### Genome-wide association studies (GWAS)

The Pan-UK Biobank Project, described in Karczewski et al.^19^, has publicly released individual GWAS in each ancestry as well as meta-analyzed GWAS across ancestries. We utilized AFR and EUR GWAS, as well as the meta-analyzed GWAS across the AFR and EUR groups, from this resource.

The phenotypes within the AoU were processed using the same strategy described in Karczewski et al.^19^, where the quantitative phenotypes were inverse-ranked normalized. We performed GWAS on the training datasets within AFR, AMR and EUR populations as described previously using the Regenie software^55^. Only the quantitative phenotypes with sample size larger than 5,000 and binary traits with case counts exceeding 100 were included for GWAS analysis. We included the following covariates: age, sex, and the first 10 PCs.

We then conducted meta-analyses of the AoU GWAS data with the UKB GWAS data, separately for EUR and AFR, as well as all ancestry groups combined. Meta-analyses across three ancestry groups within AoU were also performed. The meta-analyses were performed using the inverse-variance weighted approach in the METAL software^56^. Our analyses focused on common HapMap 3 variants only.

For additional sensitivity analyses, we downsampled the EUR, AFR, and AMR groups in AoU, as well as the EUR group in UKB, to match the smallest sample size of unrelated individuals across the training datasets for 13 quantitative traits. We excluded reticulocyte percentage in these analyses due to its small sample sizes in AoU (5,114 EUR; 2,196 AFR; and 1,479 AMR). We performed additional GWAS using data from each of the downsampled groups.

### Genetic architecture estimates

In this study, we investigated the impact of key parameters of genetic architecture on the performance of PRS. We assessed several trait-specific genetic architecture parameters, namely polygenicity (i.e. the proportion of SNPs with nonzero effects) and SNP-based heritability. To estimate polygenicity, we employed SBayesS, a summary statistics based method employing a Bayesian mixed linear model, with its default settings^57^. The input datasets for this analysis were the EUR GWAS from UKB. To estimate heritability, we conducted LD score regression analyses using LDSC^40^ based on the AoU EUR GWAS, and obtained the LDSC estimates based on the UKB EUR GWAS from Karczewski et al.^19^. We used ancestry-matched reference panels from UKB for these analyses^19^. We also estimated heritability of 13 quantitative traits (excluding reticulocyte percentage due to small sample sizes) from individual-level data using the restricted maximum likelihood (GREML) approach as implemented in GCTA^41^. We used data from the downsampled EUR, AFR, and AMR groups in AoU, as well as the EUR group in UKB. The raw phenotype values and covariates (age, sex, and the first 10 PCs) were input to GCTA-GREML.

### Genetic correlation estimates

To estimate *r_g_* between the EUR GWAS from AoU and UKB, we used the heritability Z-scores obtained from LDSC computations of heritability from AoU GWAS and as reported in Karczewski et al.^19^ from UKB GWAS. To estimate cross-ancestry *r_g_* between the EUR and AFR GWAS from AoU, and EUR and AMR GWAS from AoU, we used Popcorn^43^ based on 1000 Genomes reference panels.

### PRS construction and evaluation

We constructed PRS using three different methods: the classic pruning and thresholding (P+T) method, and two Bayesian genome-wide methods, namely PRS-CS^58^ and PRS-CSx^9^. P+T was performed using a LD *r*^2^ threshold of 0.1 and a series of *p*-value thresholds (5e-8, 5e-07, 5e-06, 5e-05, 5e-04, 5e-03, 0.05, 0.1, 1). We used the auto model, which automatically estimates the global shrinkage parameter, implemented in PRS-CS and PRS-CSx. We used ancestry-specific AoU GWAS as inputs for the three methods. For P+T and PRS-CS, multi-ancestry meta-analyzed GWAS were additionally included. In order to comprehensively explore the advantages of incorporating AoU data, we constructed PRS using UKB GWAS data independently, as well as the meta-analyzed AoU and UKB GWAS data.

The LD reference panel used was dependent on the ancestry composition of the discovery GWAS. We used LD panels that matched the respective ancestral population for ancestry-specific GWAS. Since the multi-ancestry meta-analyzed GWAS primarily comprised European individuals, we used a European-based panel, as our previous studies demonstrated that it can adequately approximate the LD structure^13,25^. We used the pre-computed LD matrices obtained from Karczewski et al.^19^ for P+T. Additionally, for PRS-CS and PRS-CSx, we employed the LD matrices provided by the software, which were computed from UKB data. We evaluated PRS performance in independent target datasets of AFR, AMR, EUR, EAS, and CSA ancestries within the AoU dataset. To evaluate the PRS performance for quantitative phenotypes, we estimated incremental R2 by accounting for the covariates: age, sex, and the first 10 PCs. Specifically, we compared two models: 1) the baseline model (phenotype ∼ covariates) and 2) the full model including PRS (phenotype ∼ PRS + covariates). Incremental *R^2^* represents the improvement in model accuracy with the inclusion of PRS. For binary phenotypes, we reported the Area Under the Receiver Operating Characteristic Curve (AUC) of PRS solely, Nagelkerker’s *R^2^*, and *R^2^* on the liability scale. In the latter case, we approximated the disease prevalence using the population prevalence. We calculated the corresponding 95% confidence intervals (CIs) of each estimate using 1,000 bootstrap iterations. Additionally, we computed odds ratios for each PRS quantile, using the first quantile as the reference. For the P+T method, we adopted a two-step evaluation approach. First, we partitioned the target datasets evenly into a validation cohort and a test cohort. Next, we fine-tuned the *p*-value threshold using the validation cohort to optimize performance. Subsequently, we evaluated the PRS performance on the test cohort using the fine-tuned *p*-value threshold. This procedure ensured a robust evaluation of the PRS performance based on the optimal thresholds. We computed relative accuracies of PRS derived from P+T with genome-wide significant SNPs vs. PRS derived from PRS-CS by dividing incremental R2 from each PRS by the incremental R^2^ from PRS_AoU-EUR_.

In an additional sensitivity analysis, we stratified all target individuals into quintiles based on their proportion of European ancestry, and evaluated PRS accuracy for quantitative traits within each quintile. The proportion of ancestries for each individual were inferred using SCOPE^60^.

### Estimates of population genetic differentiation

To characterize the genetic distance between populations across the biobanks, we measured population genetic differentiation with Wright’s fixation index, *F_st_*, computed using the “wc” method in PLINK 2.0^54^. The analyses were performed using 168,899 pruned variants.

### Individual PRS accuracy

#### Posterior effect size calculation

We used the EUR GWAS and multi-ancestry meta-analysis from AoU as inputs for PRS-CS. Using the default setting of PRS-CS, which involves 1000 MCMC (Markov Chain Monte Carlo) iterations, 500 burn-in iterations, and a thinning factor of 5, we obtained an output of 100 sets of posterior effect estimates for each variant in an *M*x100 matrix, where *M* is the number of SNPs. This matches the output based on LDPred2 in Ding et al.^39^

#### PRS accuracy

We used the “--score” flag in PLINK 2.0 to compute 100 sets of PRS for the individuals in the AMR, AFR, EUR, CSA, and EAS target groups in AoU, as well as for individuals in AoU who were not assigned to any ancestry group with the PCA approach and thus were not included in the discovery or target groups for PRS evaluation using population-level metrics. The output matrix has shape 100 × 22,703 and each cell is denoted as 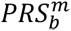, where *m* ∈ [1,22703] denotes the *m* individual and *b* ∈ [1,100] denotes the *b* set of PRS. Based on Ding et al.^39^, the individual PRS uncertainty for individual *m* for empirical analyses is calculated as *var*(*PRS^m^*), that is the variance of 100 sets of PRS. The PRS accuracy for individual *m* is defined as 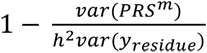, where *h*^2^ denotes the estimated heritability from LDSC and *var*(*y_residue_*) denotes the variance of residue phenotype in training data after regressing out age, sex, and the first 10 PCs. The PRS accuracies for four blood panel traits (neutrophil count, white blood cell count, mean corpuscular volume, and mean corpuscular hemoglobin), for which PRS_AoU-Multi_ performed best, and two additional polygenic traits (height and BMI) for comparison were scaled using min-max normalization ranging from 0 to 1, where the minimum and maximum values correspond to the smallest and largest PRS accuracy observed among individuals across all six traits, respectively. The correlation coefficient *R* was measured by Pearson correlation.

#### Genetic distance

We used the same strategy as described in Ding et al.^39^ to calculate genetic distance between each individual and the discovery population. Briefly, we calculated the Euclidean distance of the PCs of the individuals in the target groups from the center of the discovery data, i.e. either the EUR or all groups in AoU.

## Acknowledgements

We acknowledge helpful comments from Mark Daly and Konrad Karczewski. A.R.M is funded by the K99/R00MH117229. K.T. is funded by F31HL167378 and supported by the ECOR Claflin Award to A.R.M. A.R.M. and Y.W. are funded by U01HG011719. Additional support for this work to A.R.M. and Y.W. also comes from the European Union’s Horizon 2020 research and innovation program under grant agreement 101016775 (INTERVENEConsortium). B.P. is supported by U01HG011715. We acknowledge the contribution of All of Us participants and the program to this work.

