## Supplementary_Figures for "All of Us diversity and scale improve polygenic prediction contextually with greatest improvements for under-represented populations"

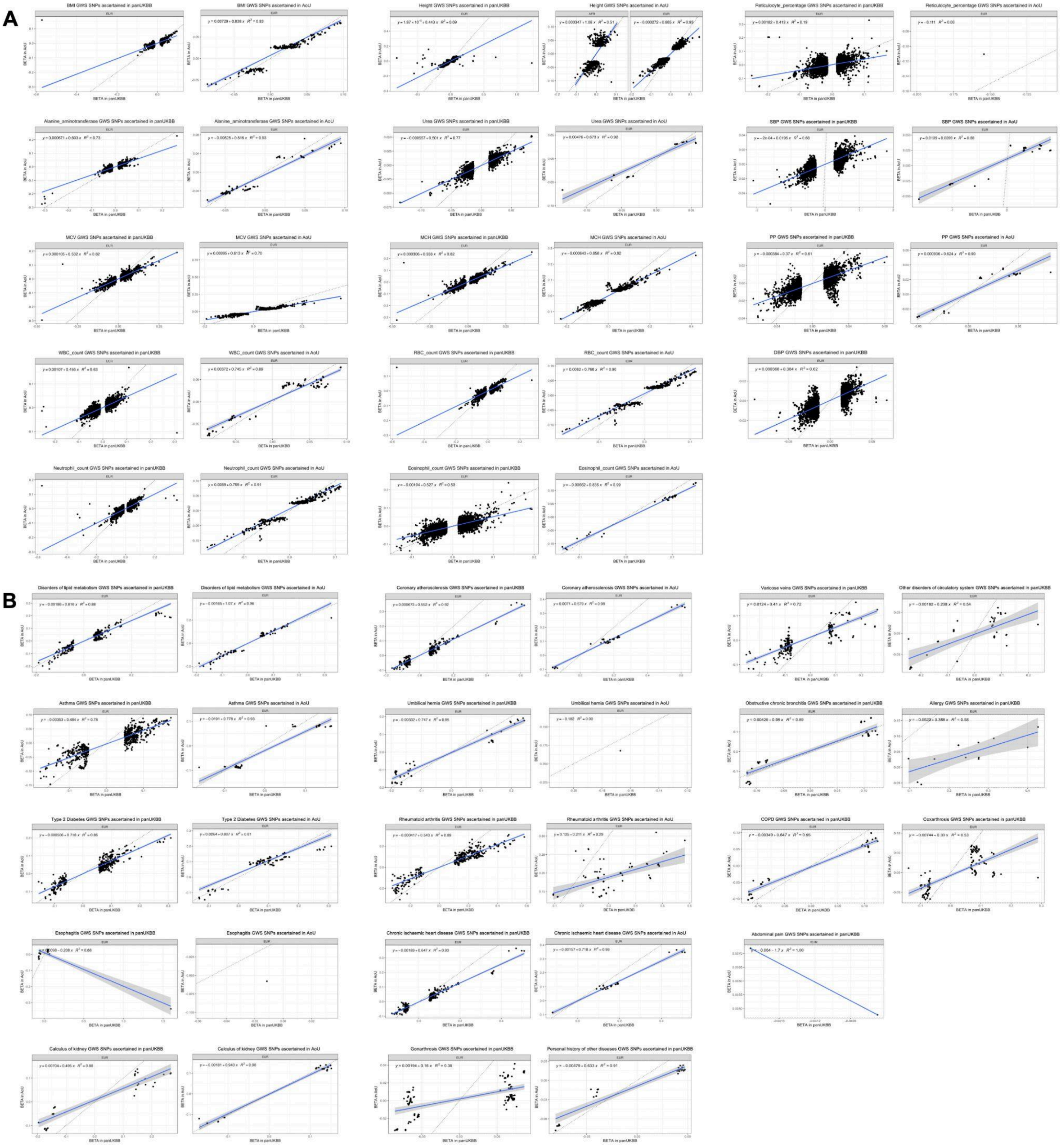


**Supplementary Figure 1. Effect size comparisons.** Effect sizes of genome-wide significant (GWS), clumped variants ascertained in AoU and UKB GWAS. Effect sizes from UKB GWAS are represented on x-axes; effect sizes from AoU GWAS are represented on y-axes. A) Quantitative phenotypes. B) Binary phenotypes.

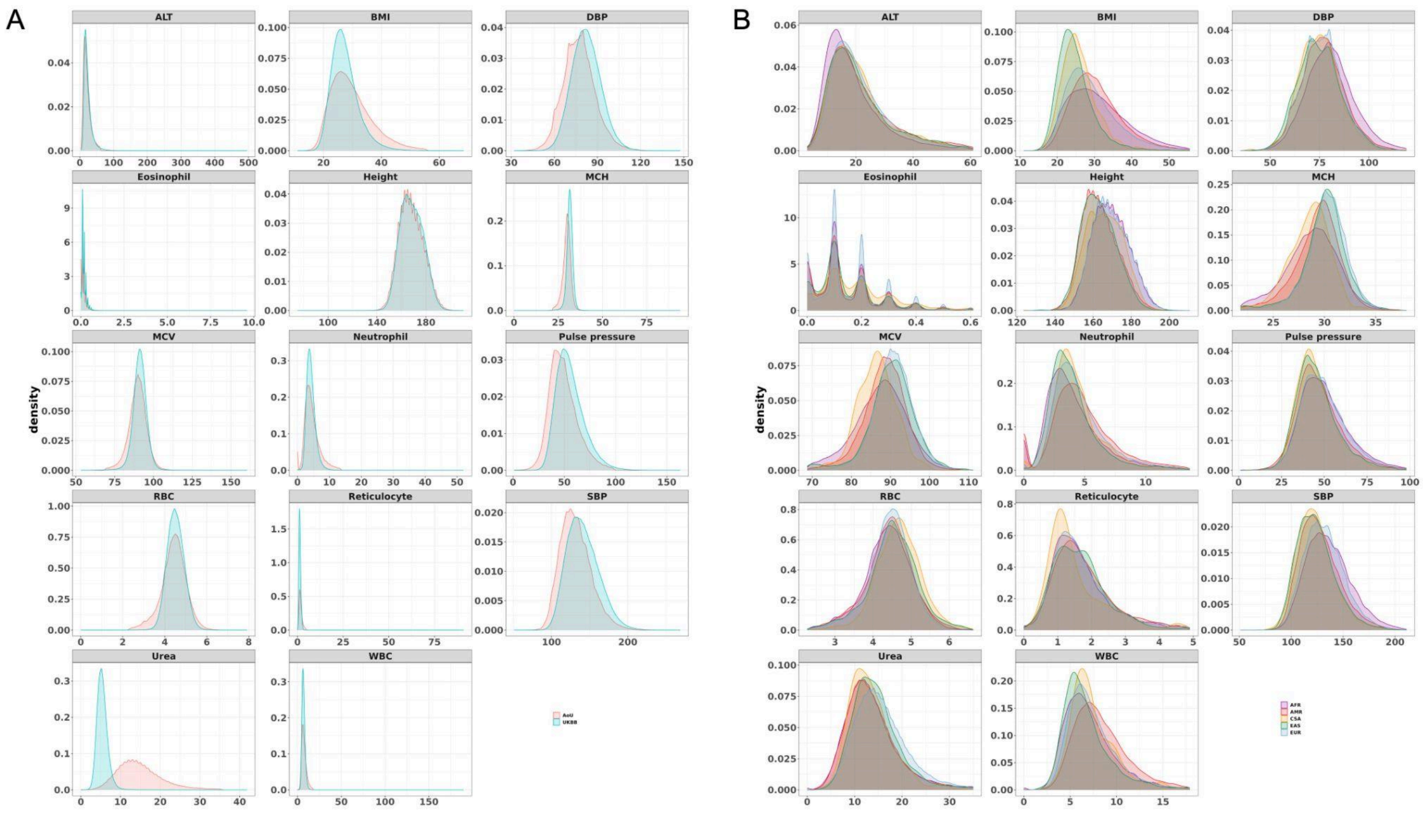


**Supplementary Figure 2. Phenotype distributions of quantitative traits in AoU vs. UKB.** Phenotype values are represented on x-axes. A) Blue curves represent distribution in UKB (EUR) and red curves represent distribution in AoU (EUR). Urea was measured with different units in UKB and AoU. B) Distributions are colored by ancestry group within AoU. (ALT = Alanine aminotransferase; BMI = body mass index; DBP = diastolic blood pressure; Eosinophil = eosinophil count; MCH = mean corpuscular hemoglobin; MCV = mean corpuscular volume; Neutrophil = neutrophil count; RBC = red blood cell count; Reticulocyte = reticulocyte percentage; SBP = systolic blood pressure; WBC = white blood cell count)

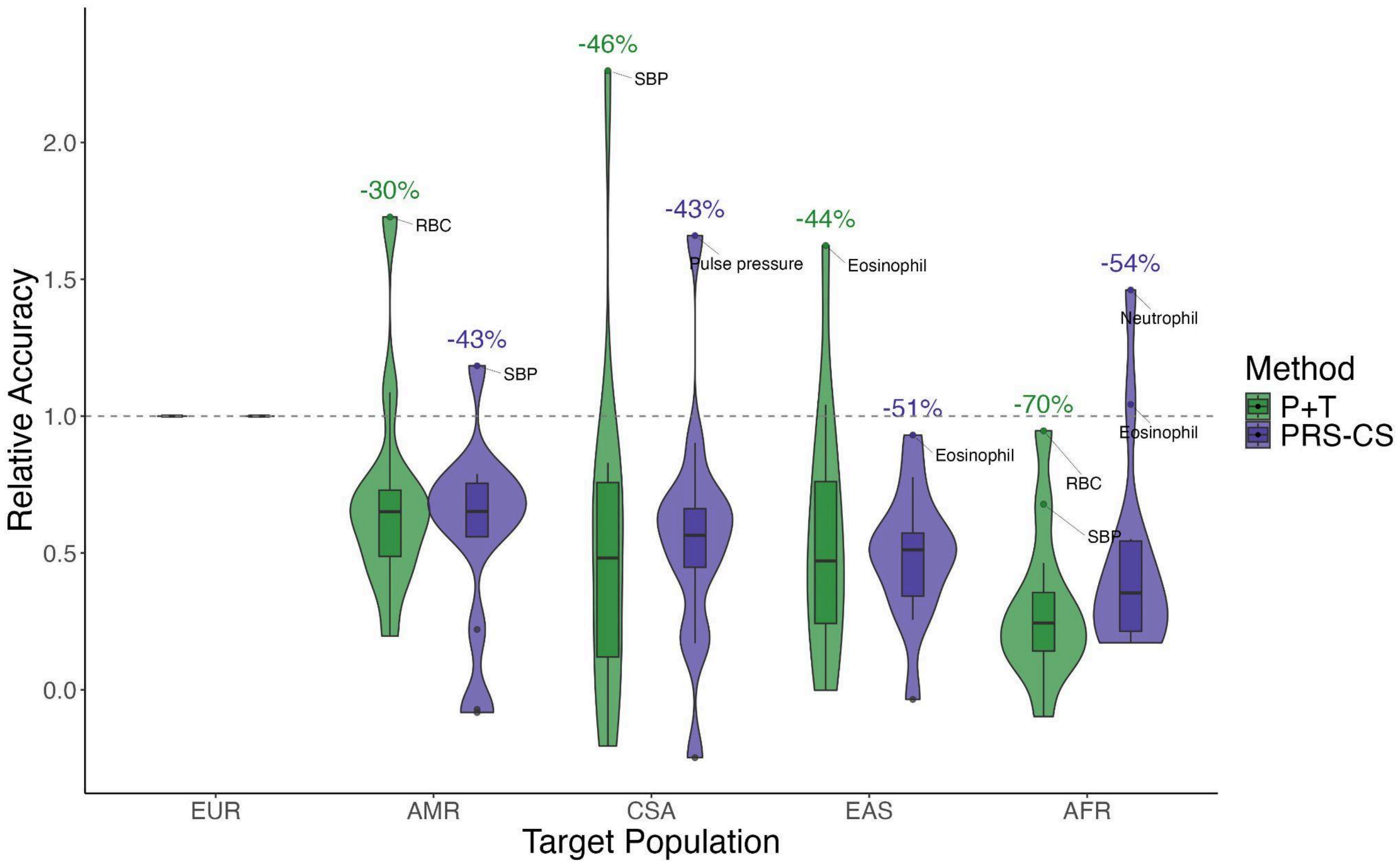


**Supplementary Figure 3. Relative accuracy of PRS derived from P+T vs. PRS-CS.** PRS were derived from AoU EUR GWAS and evaluated in target populations in AoU, as shown on x-axis. Relative accuracy was computed as the performance of $PRS_{AoU\text{-}EUR}$ in the target population / $PRS_{AoU\text{-}EUR}$ in the target EUR population. 13 quantitative traits (reticulocyte percentage was excluded because of limited sample sizes) are represented in this plot; phenotypes with relative accuracies exceeding the third quartile of the relative accuracy distribution by more than 1.5 times the interquartile range (IQR) are labeled as outliers. Average decays in accuracy across the phenotypes per method and target population are also noted.

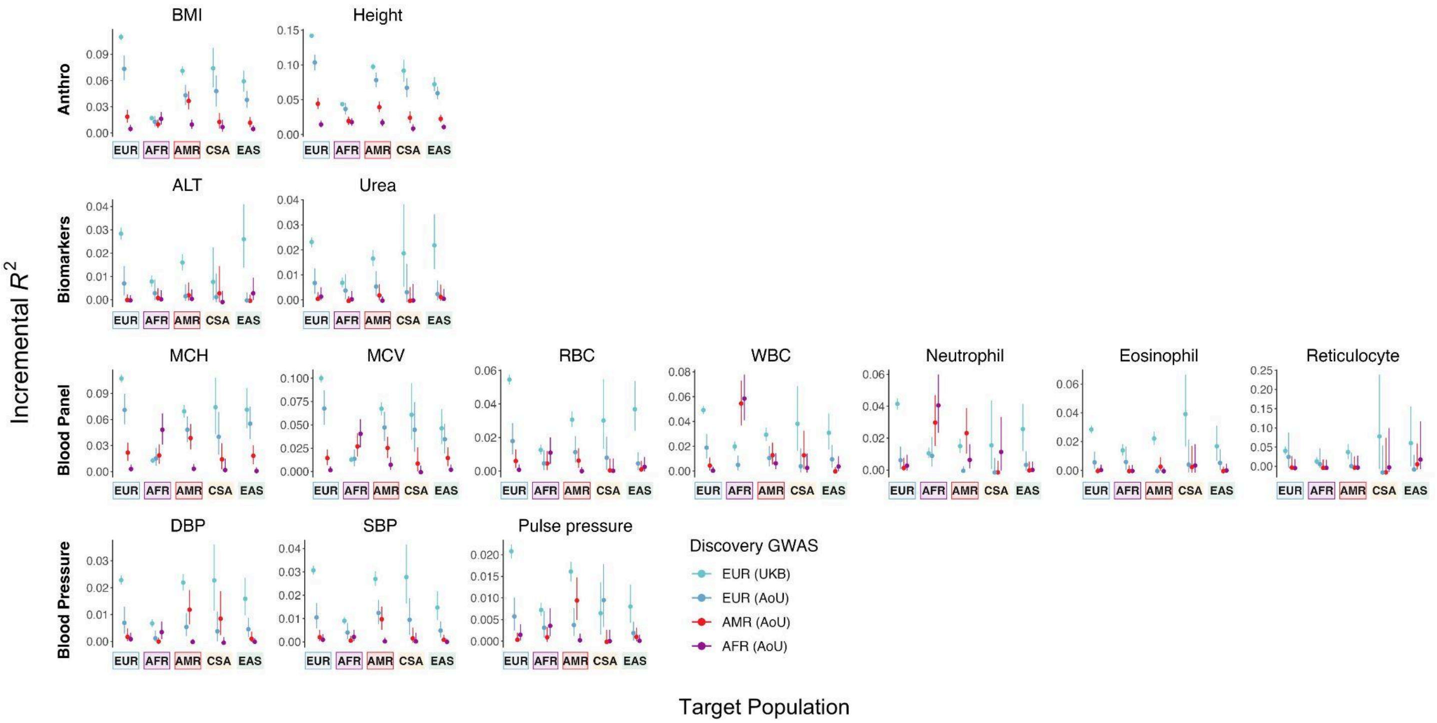


**Supplementary Figure 4. Performance of PRS derived from single-ancestry discovery GWAS from AoU and UKB for quantitative traits.** Quantitative traits are divided into categories: anthropometric traits, biomarkers, blood panel traits, and blood pressure traits. PRS were constructed from PRS-CS applied to EUR GWAS from UKB and AoU, as well as AMR and AFR GWAS from AoU. Target populations from AoU are shown on x-axes and target populations with ancestry-matched PRS are outlined; PRS performance as measured by incremental $R^2$ is shown on y-axes.

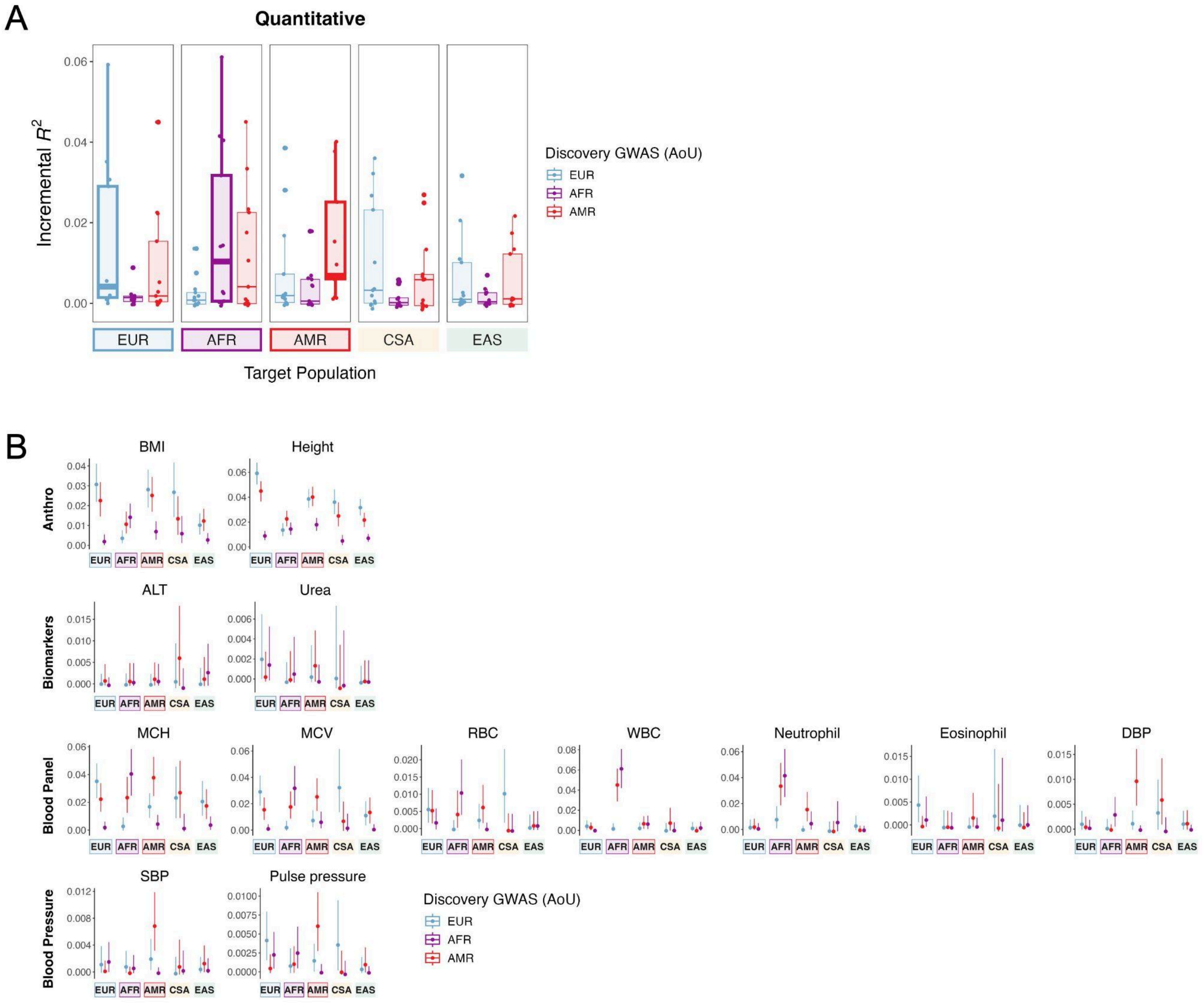


**Supplementary Figure 5. Performance of PRS derived from downsampled single-ancestry discovery GWAS from AoU for quantitative traits.** A) Each point represents a phenotype, with PRS constructed from PRS-CS applied to EUR, AMR and AFR GWAS from AoU with matching sample sizes reported here. Target populations with ancestry-matched PRS are outlined. B) Quantitative traits are divided into categories: anthropometric traits, biomarkers, blood panel traits, and blood pressure traits. PRS were constructed from PRS-CS applied to EUR, AMR and AFR GWAS from AoU with matching sample sizes. Reticulocyte percentage was excluded in these analyses because of limited sample sizes in AoU. Target populations from AoU are shown on x-axes and target populations with ancestry-matched PRS are outlined; PRS performance as measured by incremental $R^2$ is shown on y-axes.

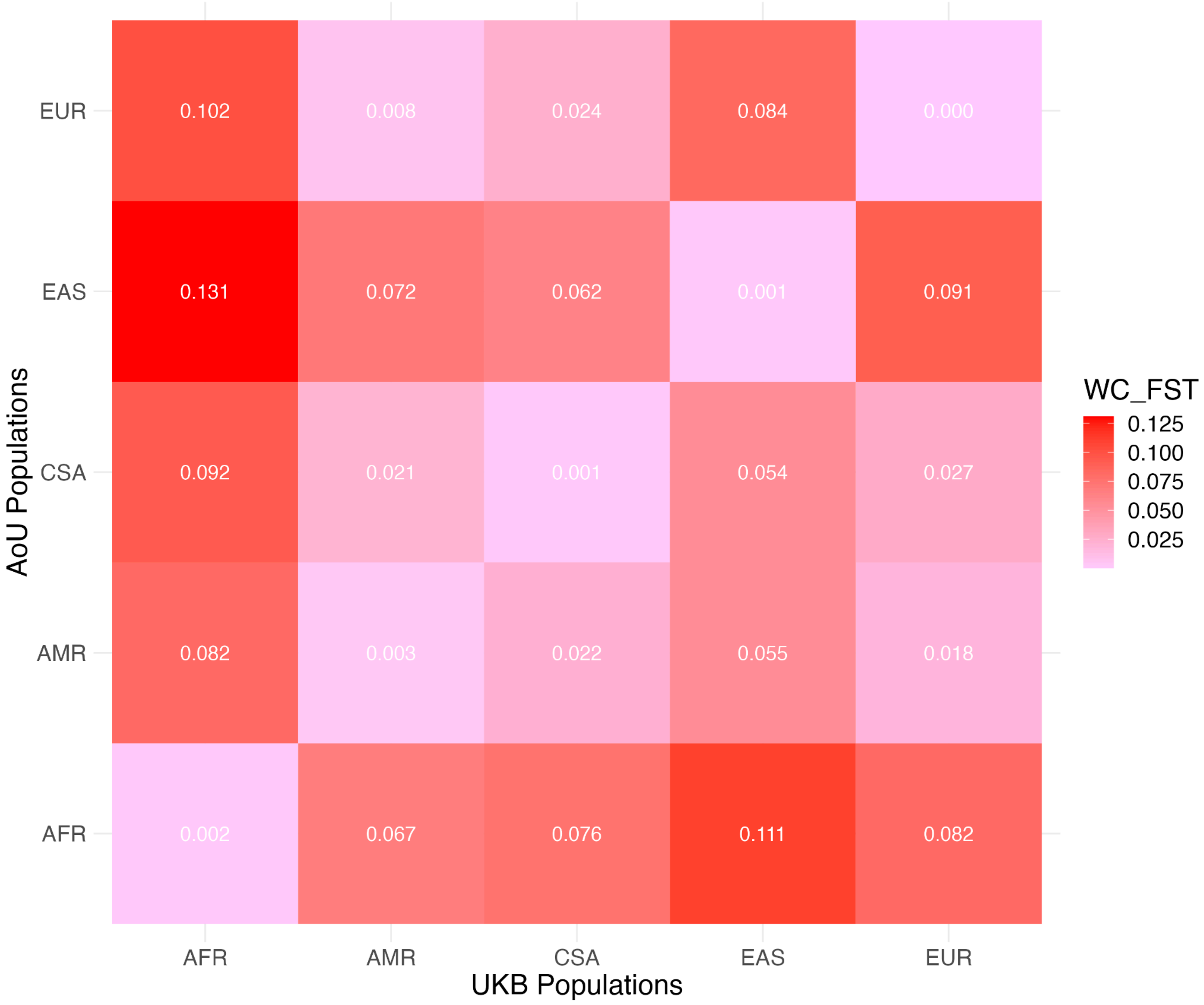


**Supplementary Figure 6. Genetic distance between populations in AoU and UKB.** Population genetic differentiation, as measured by Wright's fixation index, $F_{st}$, between genetic ancestry groups in UKB vs. AoU.

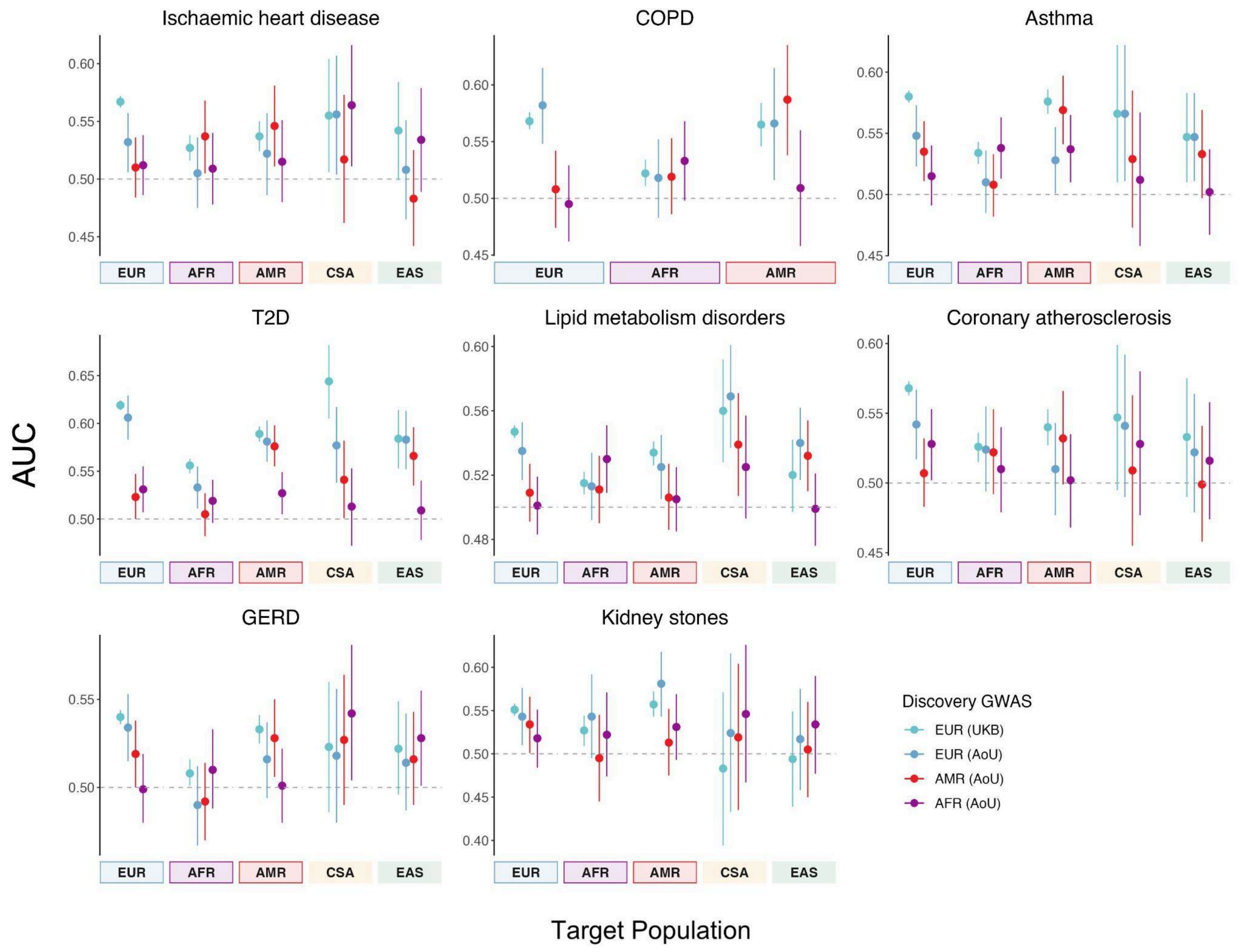


**Supplementary Figure 7. Performance of PRS derived from single-ancestry discovery GWAS from AoU and UKB for binary phenotypes.** PRS were constructed from PRS-CS applied to EUR GWAS from UKB and AoU, as well as AMR and AFR GWAS from AoU. Target populations from AoU are shown on x-axes and target populations with ancestry-matched PRS are outlined; PRS performance as measured by AUC is shown on y-axes.

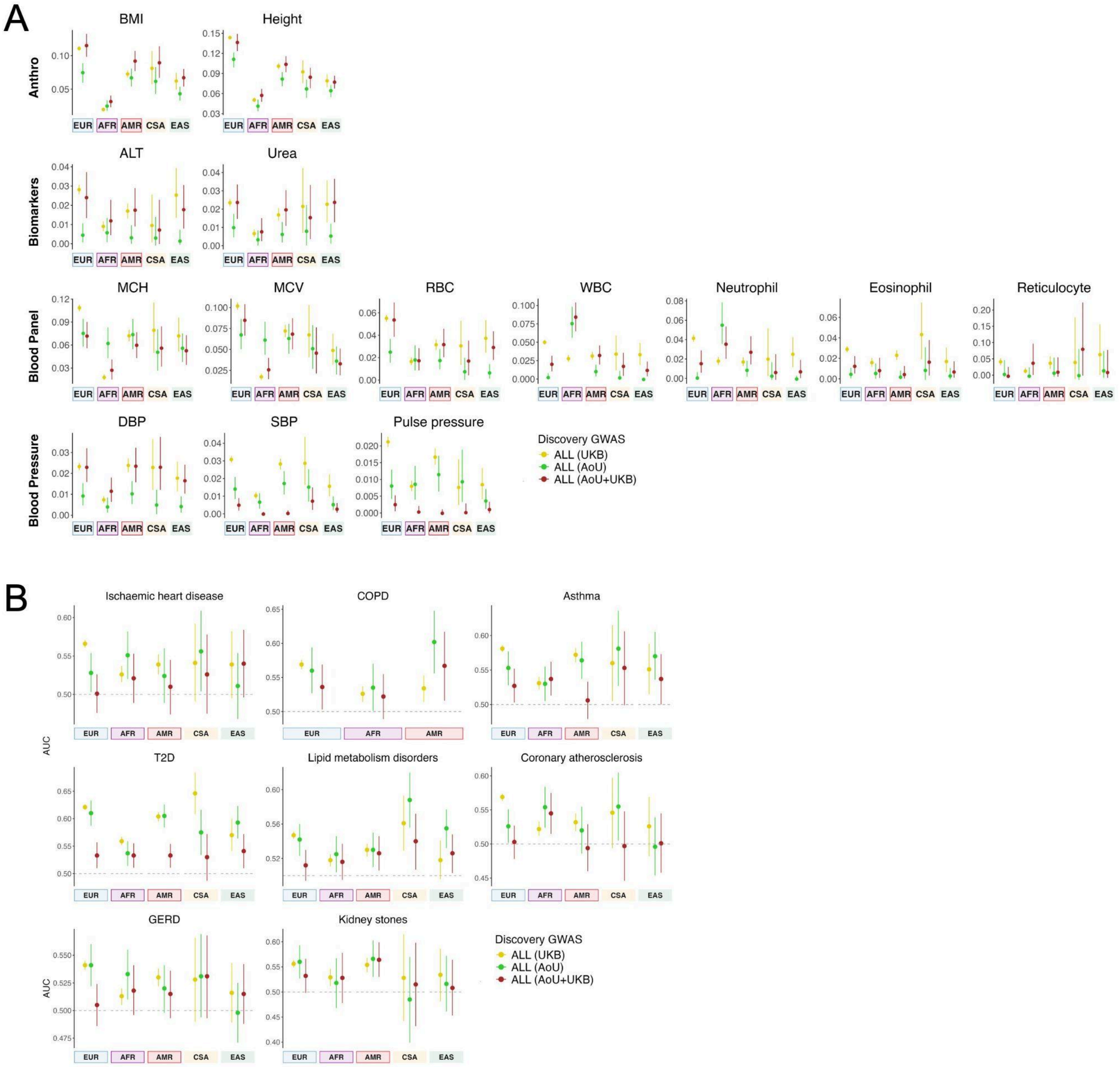


**Supplementary Figure 8. Performance of PRS derived from multi-ancestry discovery GWAS from AoU and UKB.** PRS were constructed from PRS-CS applied to multi-ancestry GWAS from UKB and AoU, as well as cross-biobank, multi-ancestry meta-analyses. Target populations from AoU are shown on x-axes and target populations with ancestry-matched PRS are outlined. A) Quantitative traits are divided into categories: anthropometric traits, biomarkers, blood panel traits, and blood pressure traits. PRS performance as measured by incremental $R^2$ is shown on y-axes. B) PRS performance for binary traits as measured by AUC is shown on y-axes.

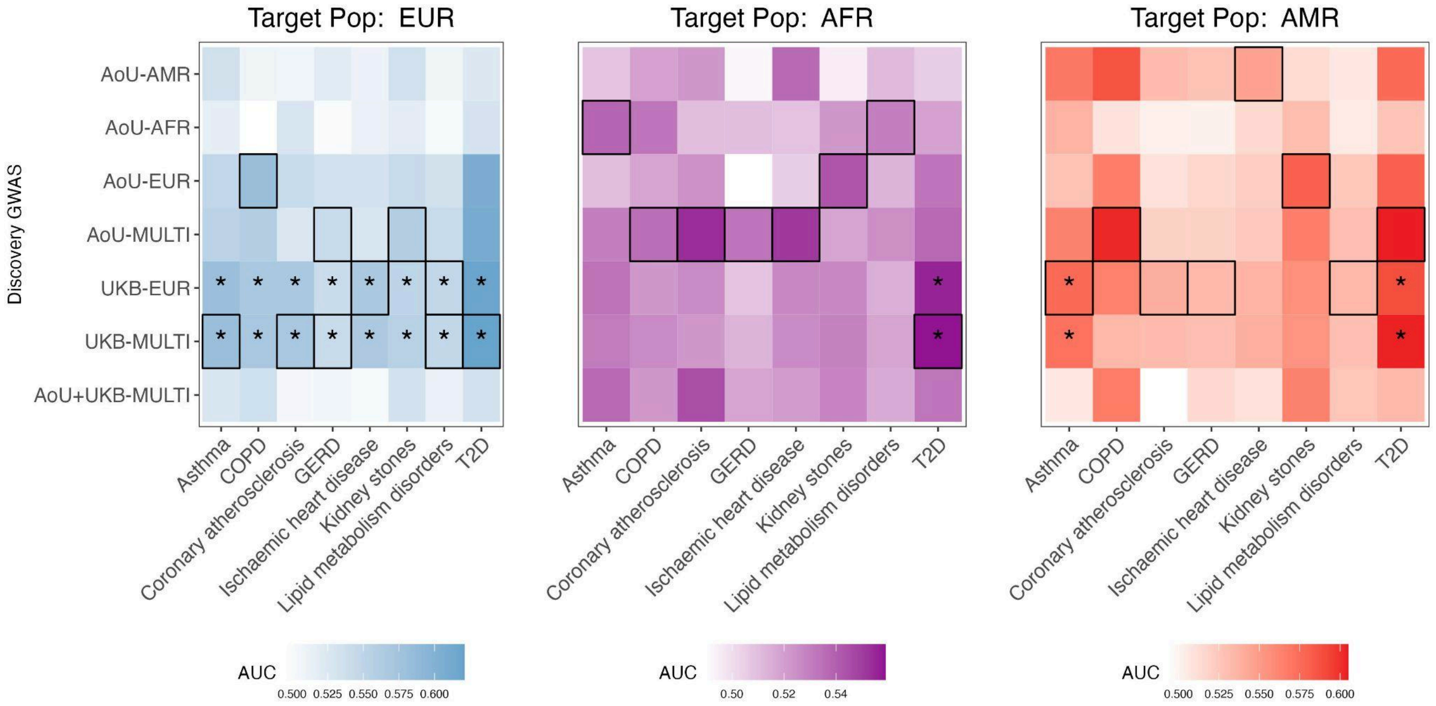


**Supplementary Figure 9. Performance of all PRS models for binary phenotypes.** PRS derived from discovery GWAS, denoted on y-axis, for binary phenotypes denoted on x-axis. PRS model with highest AUC per trait is outlined. Asterisk indicates AUC significantly greater than 0.5 (t-test, $p < 0.05$).

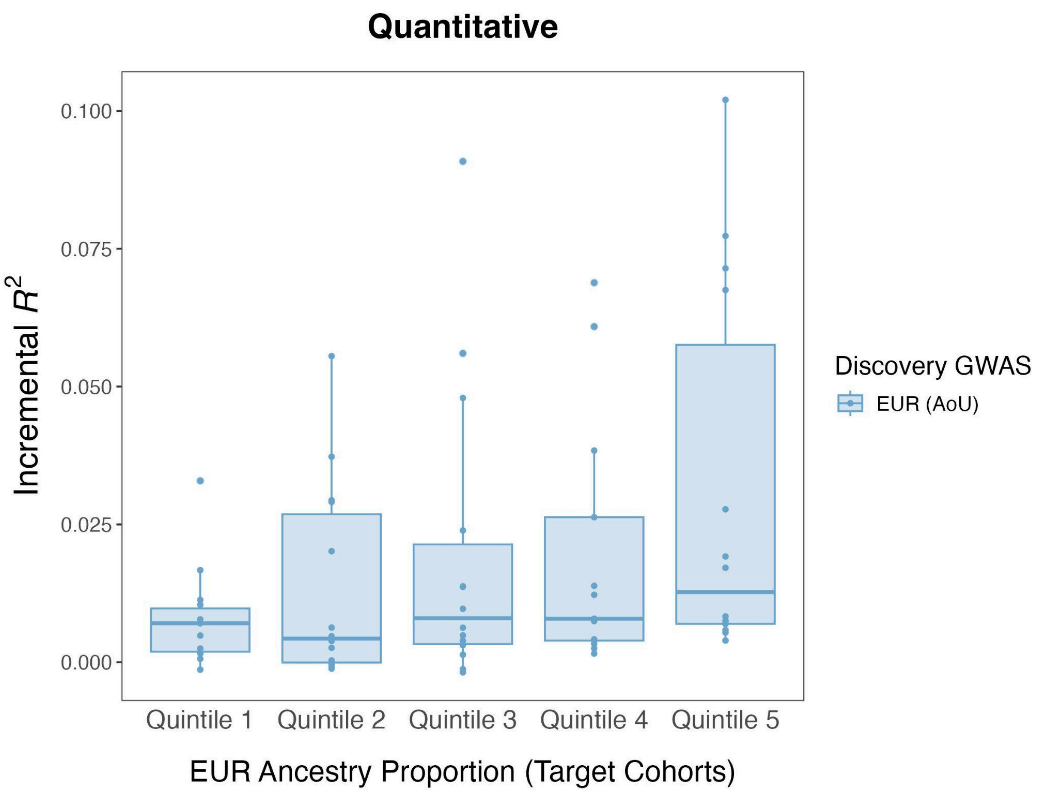


**Supplementary Figure 10. Performance of PRS for quantitative traits across target cohorts stratified by proportion of European ancestry.** Individuals across all target cohorts in AoU were stratified by proportion of European ancestry, shown on x-axis. Quintile 1 indicates smallest proportion of European ancestry; quintile 5 indicates largest proportion of European ancestry. Performance of PRS from AoU EUR GWAS, as measured by incremental $R^2$, is shown on y-axis. Each point represents a phenotype.

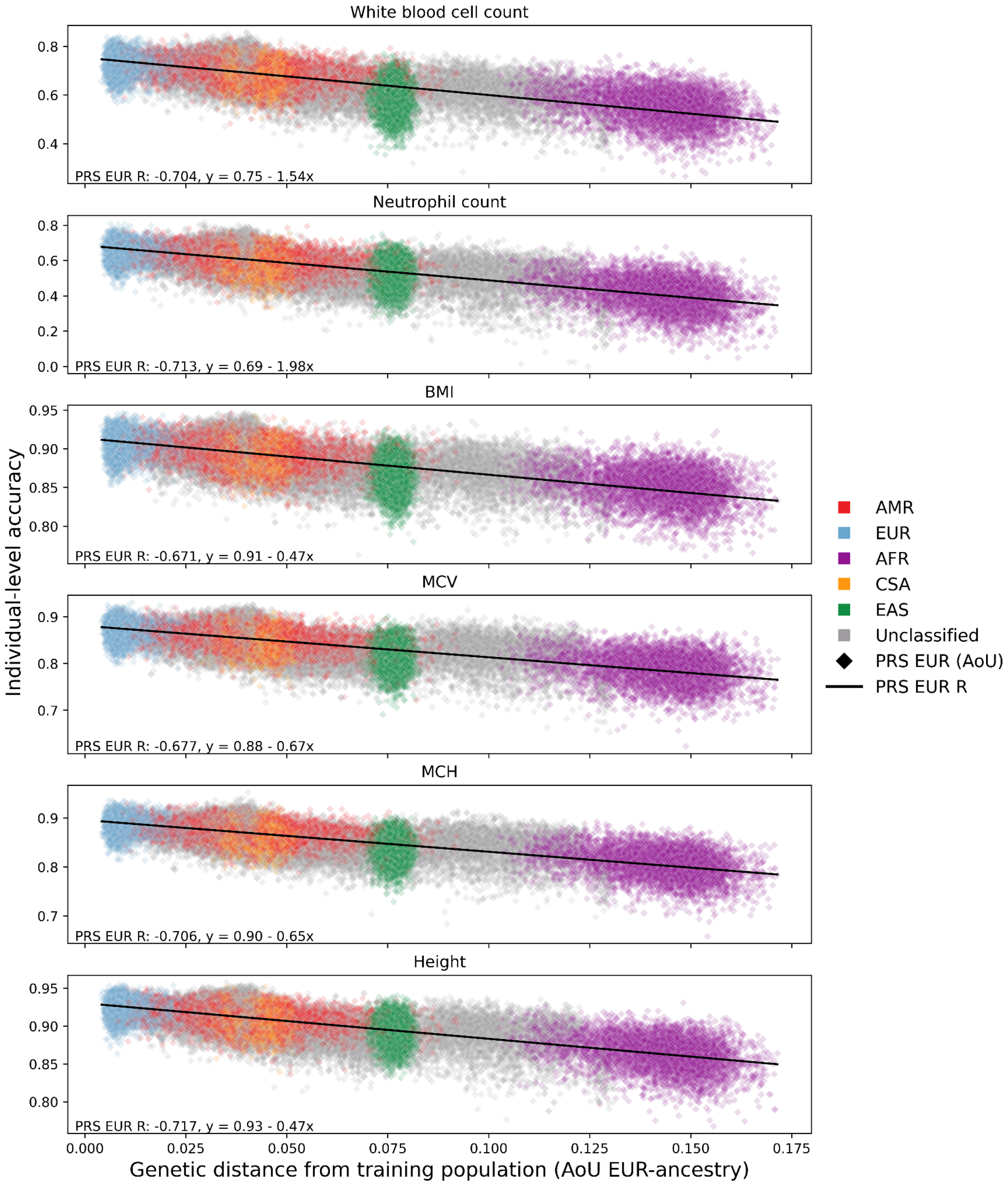


**Supplementary Figure 11. Individual-level accuracy for PRS derived from AoU EUR GWAS.** Each point represents a target individual in AoU. The x-axis represents the genetic distance (GD) of each target individual from the EUR discovery group in AoU. The y-axis shows the PRS accuracy, which was scaled to enable cross-trait comparisons of decay in accuracy as a function of GD; as a result, proportions of genetic liability explained by PRS for each individual are not represented here. R was calculated as the correlation between GD and PRS accuracy from a two-sided Pearson correlation test. The colors represent genetic ancestry groups as inferred by PCA.

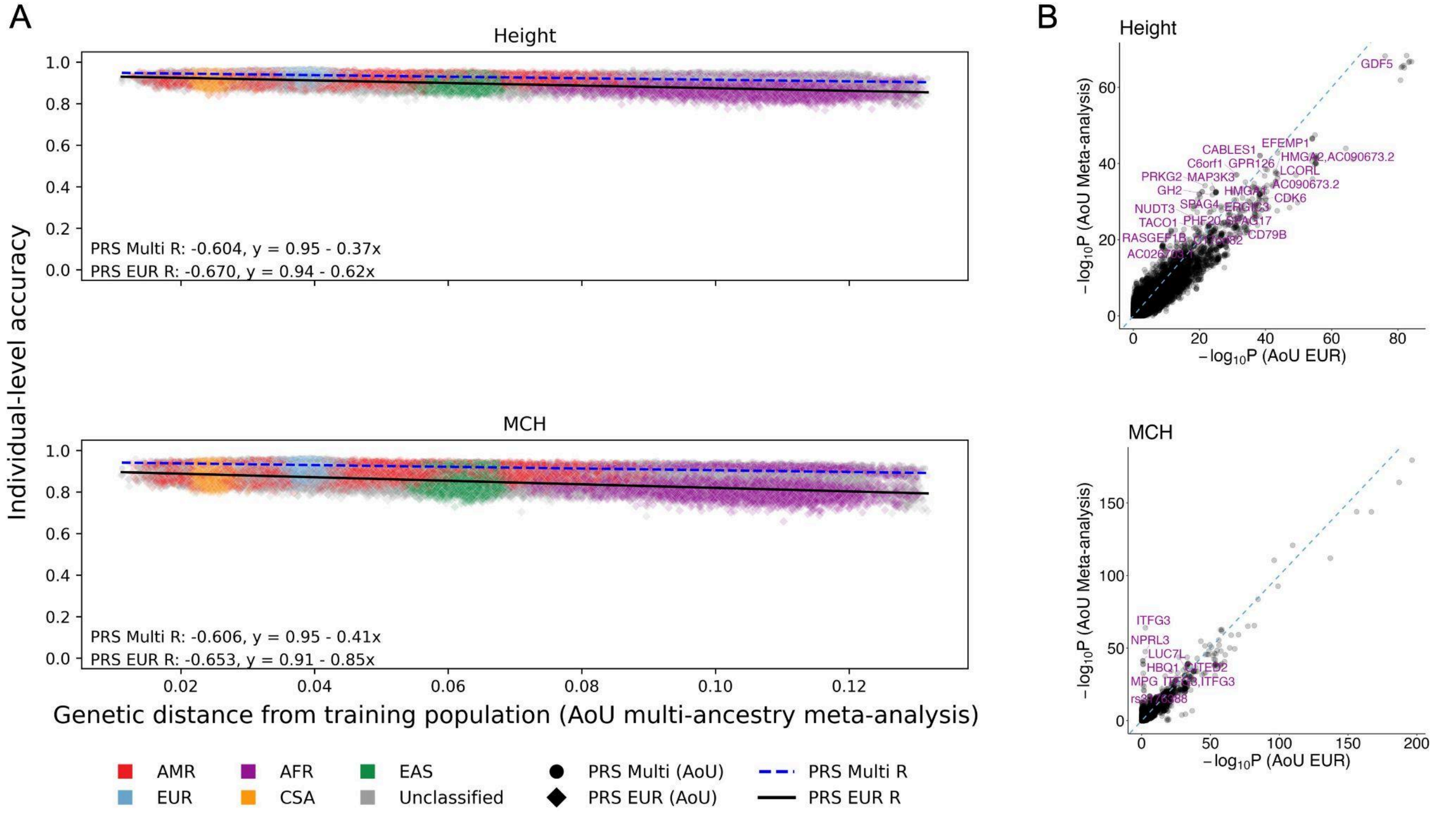


**Supplementary Figure 12**. **Individual-level accuracy for height and MCH PRS derived from AoU multi-ancestry meta-analyses.** A) Individual-level accuracy derived from AoU multi-ancestry meta-analyses and EUR GWAS across target individuals in AoU, represented by each point. The x-axis represents the genetic distance (GD) of each target individual from the combined discovery populations included in the AoU multi-ancestry meta-analyses. The y-axis shows the PRS accuracy, which was scaled to enable cross-trait comparisons of decay in accuracy as a function of GD; as a result, proportions of genetic liability explained by PRS for each individual are not represented here. R was calculated as the correlation between GD and PRS accuracy from a two-sided Pearson correlation test. The colors represent genetic ancestry groups as inferred by PCA. B) Comparison of GWAS significance in AoU multi-ancestry meta-analyses and AoU EUR GWAS for height and MCH. SNPs tested in both the AoU multi-ancestry meta-analyses and EUR GWAS are represented by each point. SNPs reaching genome-wide significance ($p < 5e-8$) in the AoU meta-analysis and AoU AFR GWAS for each phenotype are annotated. Dashed lines indicate $y=x$; x- and y-axis scales are specific to each phenotype and differ according to scale of significance in meta-analyses vs. EUR GWAS.

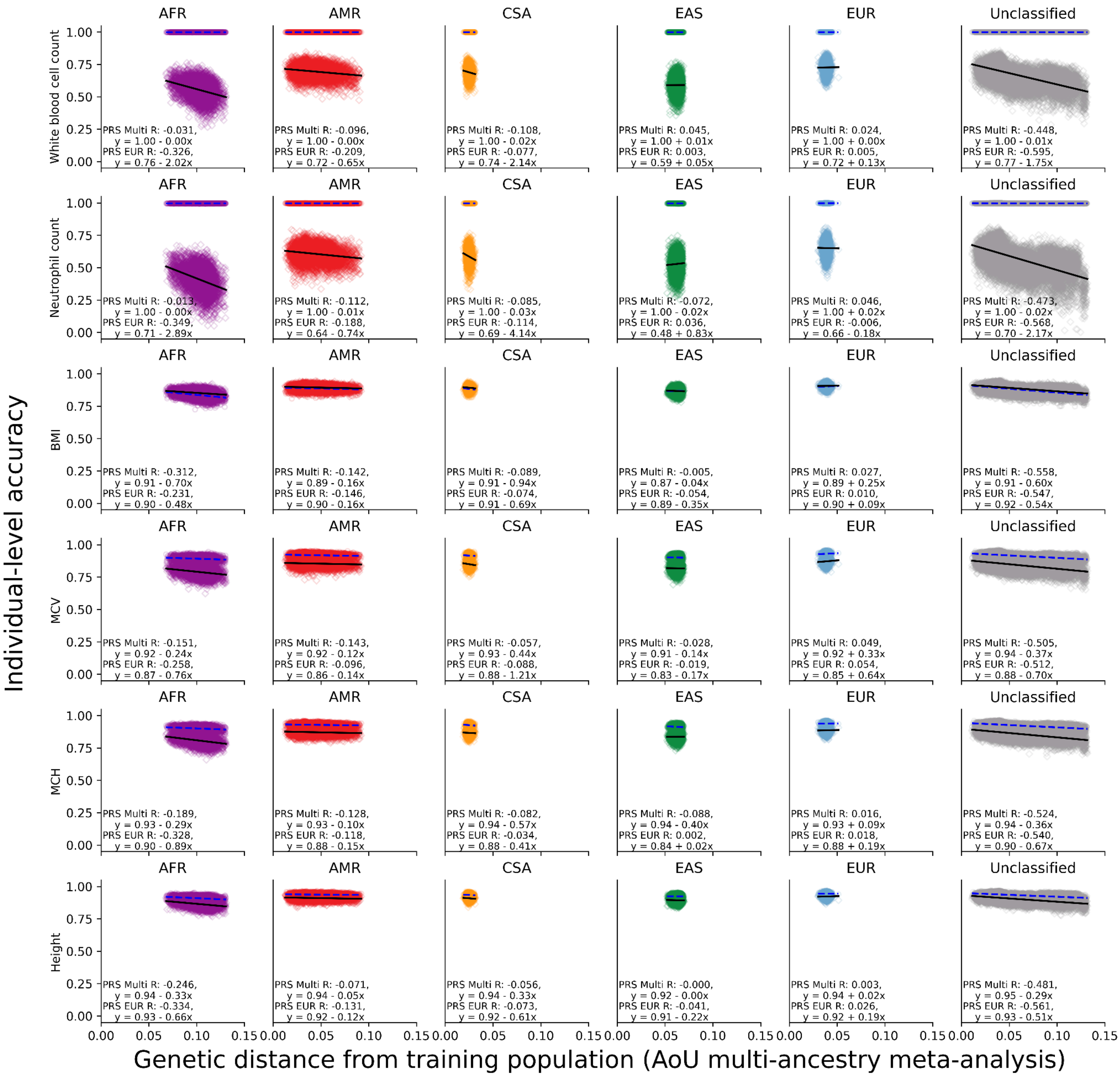


**Supplementary Figure 13. Individual-level accuracy for 6 phenotypes across the 5 target ancestry groups in AoU and individuals not assigned to an ancestry group.** Individual-level accuracy derived from AoU multi-ancestry meta-analyses and EUR GWAS across target individuals in AoU assigned to an ancestry group and individuals in AoU not assigned to any ancestry group. The x-axis represents the genetic distance (GD) of each target individual from the combined discovery populations included in the AoU multi-ancestry meta-analyses. The y-axis shows the PRS accuracy, which was scaled to enable cross-trait comparisons of decay in accuracy as a function of GD; as a result, proportions of genetic liability explained by PRS for each individual are not represented here. R was calculated as the correlation between GD and PRS accuracy from a two-sided Pearson correlation test.

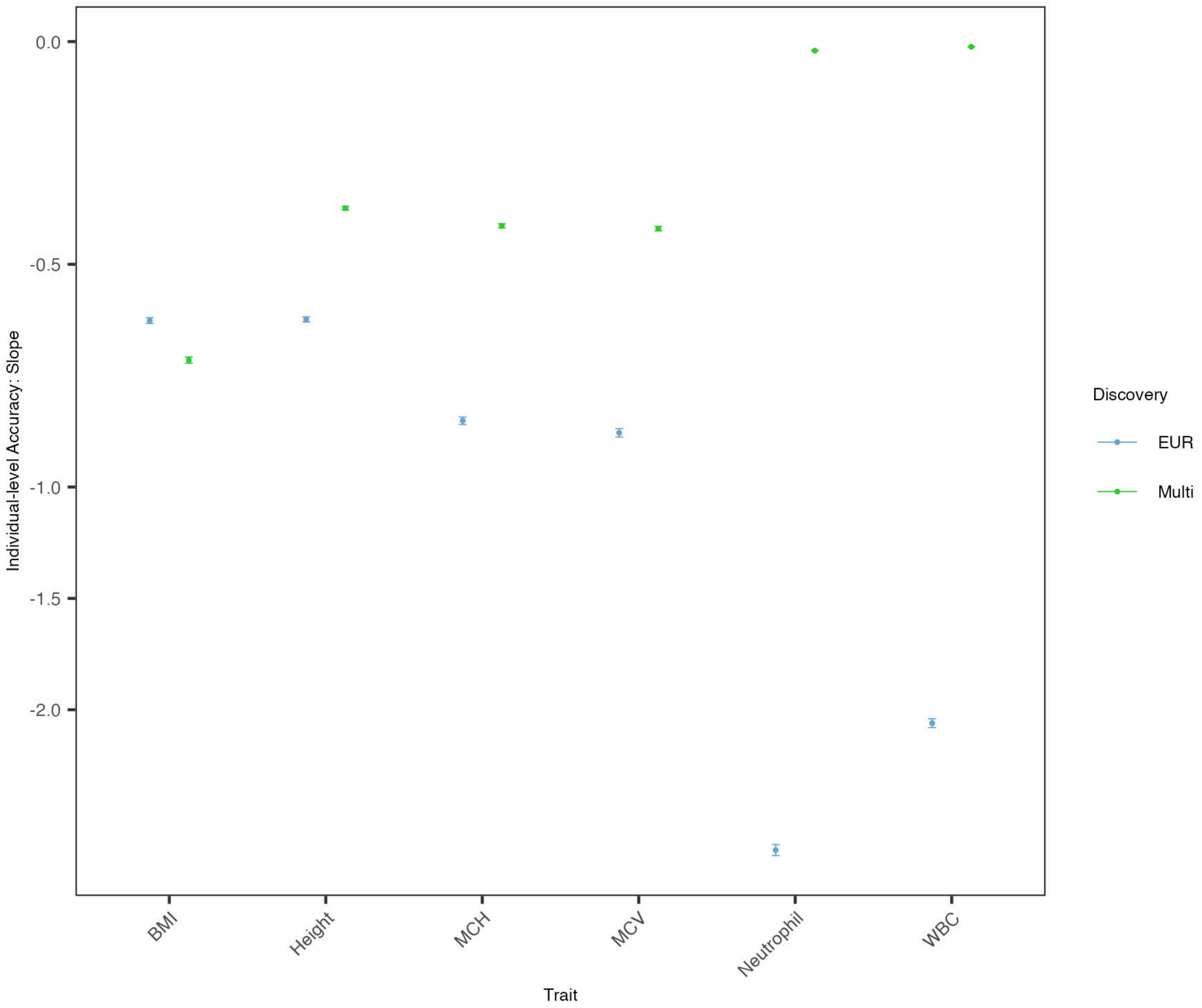


**Supplementary Figure 14. Slope estimates of individual-level PRS accuracy across target individuals for 6 quantitative traits.** Slopes represent decay in individual-level PRS accuracy as the genetic distance of target individuals from AoU discovery populations (either EUR or combined multi-ancestry populations) increases.

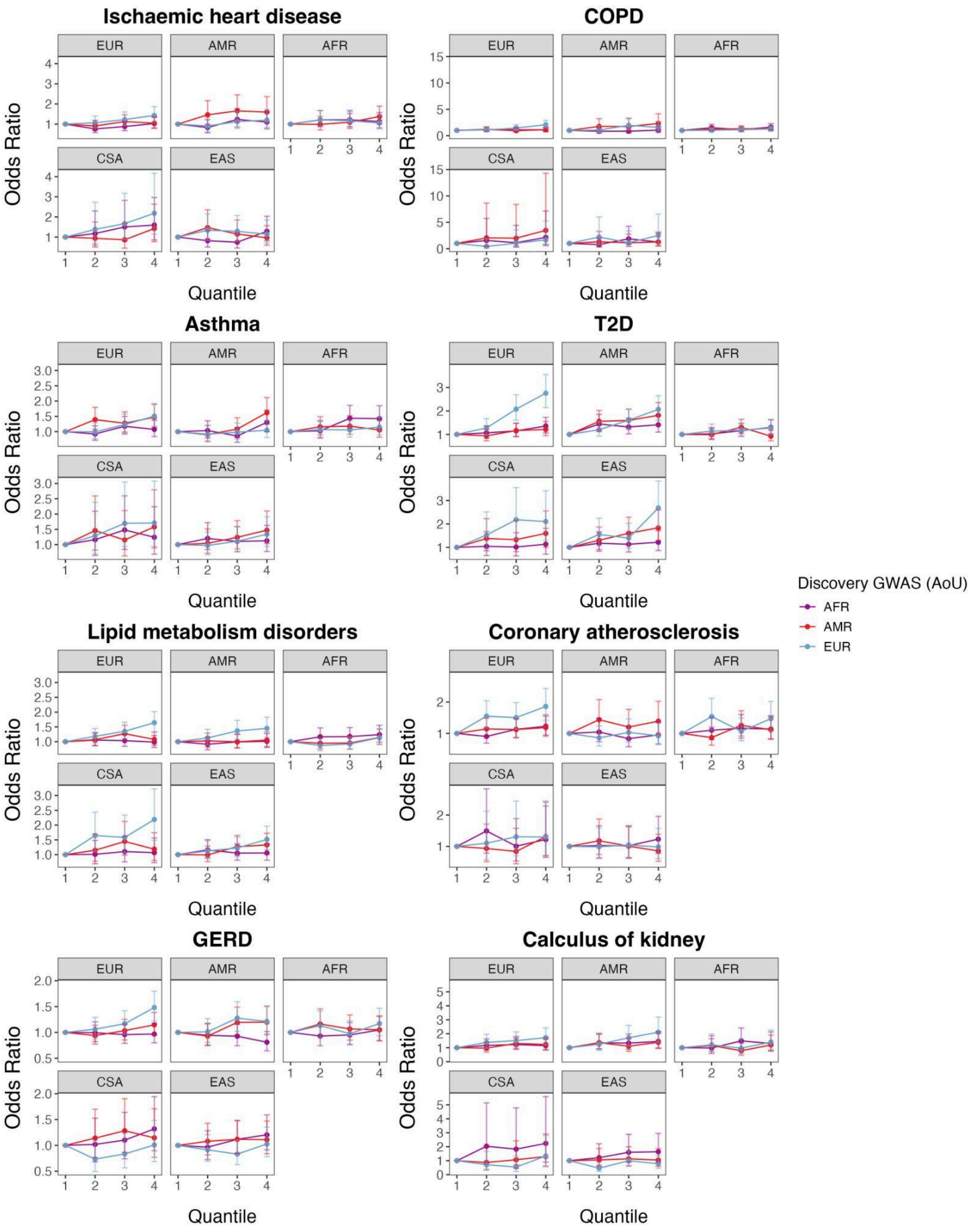


**Supplementary Figure 15. Disease risk stratification by PRS distribution.** Target populations were stratified by PRS quantile, as shown on x-axes. PRS were derived from the AoU AFR, AMR, and EUR discovery GWAS. The reference group was the first quantile.

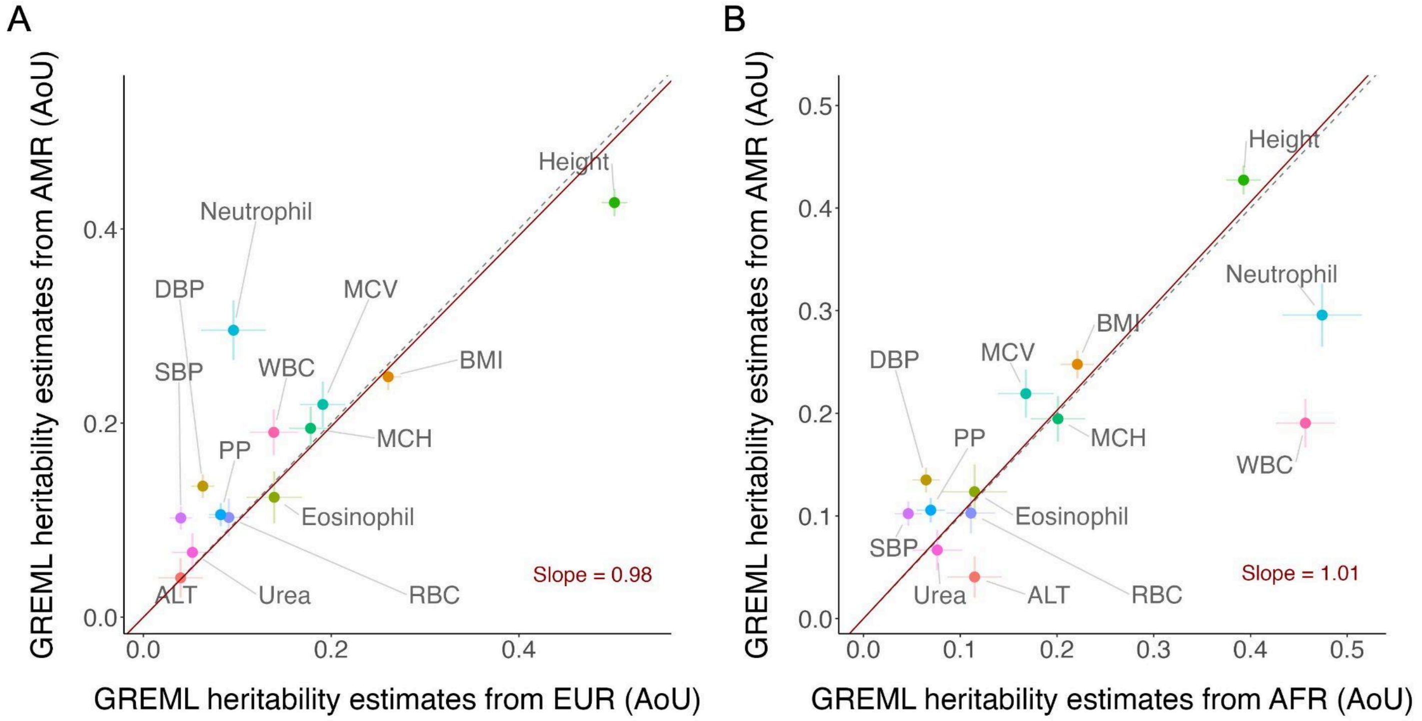


**Supplementary Figure 16. Comparison of heritability estimates across quantitative traits within populations in AoU.** Population groups in AoU were downsampled to match the smallest sample size of unrelated individuals across training datasets for each phenotype shown. GREML was applied to individual-level data from these groups to calculate SNP-based heritability estimates. Regression slopes were computed using the Deming regression method(Deming 1943). Dashed lines indicate y=x. A) Comparison of heritability estimates from EUR vs. AMR groups from AoU. B) Comparison of heritability estimates from AFR vs. AMR groups from AoU.
